# Accurate functional classification of thousands of *BRCA1* variants with saturation genome editing

**DOI:** 10.1101/294520

**Authors:** Gregory M. Findlay, Riza M. Daza, Beth Martin, Melissa D. Zhang, Anh P. Leith, Molly Gasperini, Joseph D. Janizek, Xingfan Huang, Lea M. Starita, Jay Shendure

## Abstract

**Variants of uncertain significance (VUS) fundamentally limit the utility of genetic information in a clinical setting. The challenge of VUS is epitomized by *BRCA1*, a tumor suppressor gene integral to DNA repair and genomic stability. Germline *BRCA1* loss-of-function (LOF) variants predispose women to early-onset breast and ovarian cancers. Although *BRCA1* has been sequenced in millions of women, the risk associated with most newly observed variants cannot be definitively assigned. Data sharing attenuates this problem but it is unlikely to solve it, as most newly observed variants are exceedingly rare. In lieu of genetic evidence, experimental approaches can be used to functionally characterize VUS. However, to date, functional studies of *BRCA1* VUS have been conducted in a *post hoc*, piecemeal fashion. Here we employ saturation genome editing to assay 96.5% of all possible single nucleotide variants (SNVs) in 13 exons that encode functionally critical domains of BRCA1. Our assay measures cellular fitness in a haploid human cell line whose survival is dependent on intact *BRCA1* function. The resulting function scores for nearly 4,000 SNVs are bimodally distributed and almost perfectly concordant with established assessments of pathogenicity. Sequence-function maps enhanced by parallel measurements of variant effects on mRNA levels reveal mechanisms by which loss-of-function SNVs arise. Hundreds of missense SNVs critical for protein function are identified, as well as dozens of exonic and intronic SNVs that compromise *BRCA1* function by disrupting splicing or transcript stability. We predict that these function scores will be directly useful for the clinical interpretation of cancer risk based on *BRCA1* sequencing. Furthermore, we propose that this paradigm can be extended to overcome the challenge of VUS in other genes in which genetic variation is clinically actionable.**

---

Despite our rapidly advancing knowledge of the genetic underpinnings of human disease, our ability to predict the phenotypic consequences of an arbitrary genetic variant in a human genome remains poor. This problem manifests most poignantly in the large numbers of ‘variants of uncertain significance’ (VUS) identified in ‘clinically actionable’ genes, *i.e.* genes that are already etiologically linked with a specific disease, and for which a definitive interpretation of the variant as benign or pathogenic would significantly impact clinical care^1,2^.

The gene that perhaps best highlights the challenge of VUS is *BRCA1*. Germline variants that disrupt *BRCA1* function are associated with a hereditary predisposition to breast and ovarian cancer^3–6^. Functionally disruptive germline variants in *BRCA1* are clinically actionable, *e.g.* by more aggressive screening or prophylactic surgery, interventions which lead to improved outcomes^7,8^. Furthermore, functionally disruptive somatic *BRCA1* mutations influence how tumors respond to specific therapeutic agents, *e.g.* PARP inhibitors^9–11^. Clinical sequencing of *BRCA1*, as well as many other genes linked to cancer predisposition such as *BRCA2, PALB2, BARD1, ATM*, etc., has the potential to implicate specific variants in disease^12^. Documented pathogenic *BRCA1* variants in the ClinVar database include complete or partial gene deletions, frameshifting insertions and deletions (indels), nonsense SNVs, missense variants detrimental to protein stability and function, and both intronic and exonic variants that perturb splicing^13^. However, as of January 2018, over half of *BRCA1* SNVs in ClinVar are classified as VUS. VUS are typified by rare missense SNVs, but also include variants potentially affecting mRNA production, such as SNVs near splice junctions. Further indicative of the challenge of variant interpretation, ClinVar is replete with *BRCA1* variants that have received conflicting interpretations from different experts. Of 3,936 germline *BRCA1* SNVs currently represented in ClinVar, only 983 are classified by an expert panel as ‘benign’ or ‘pathogenic’ without conflicting interpretations.

There are two major approaches for resolving VUS. The first approach, data sharing, relies on the expectation that as *BRCA1* is sequenced in increasing numbers of individuals^14^, the recurrent observation of a specific variant in multiple individuals who either have or have not developed breast and/or ovarian cancer will enable the definitive interpretation of that variant. However, although this may be possible for some variants, given that the vast majority of potential SNVs in *BRCA1* are exceedingly rare^15,16^ and that the phenotype is incompletely penetrant, it may be decades or centuries before sufficient numbers of humans are included in genotype-phenotype studies to accurately quantify cancer risk for each individual rare variant.

The second approach, functional assessment, has spurred the development of diverse *in vitro* assays for *BRCA1*^17^. As the homology-directed DNA repair (HDR) function of BRCA1 is key for tumor suppression, one commonly used assay involves expressing a BRCA1 variant in cells and assessing the integrity of the cells’ HDR pathway via inducing repair of a double strand DNA break in a fluorescent reporter construct^18,19^. Other approaches include assays for embryonic stem cell viability^20^, cell sensitivity to chemotherapeutic drugs^20^, binding to known partners such as BARD1^18,21^, and minigene-based splicing assays^22,23^. Computational tools can predict variant effects based on features such as amino acid conservation. However, although many such metrics correlate with pathogenicity, at present no computational tool is sufficiently accurate to be used for the clinical interpretation of newly observed *BRCA1* variants in the absence of genetic or experimental evidence^24,25^.

Functional assessment of *BRCA1* variants has historically been limited in several ways. Chiefly, experimental studies are *post hoc* and have not kept pace with the scaling of *BRCA1* sequencing and the accumulation of VUS. Additionally, assays that express variants as cDNA-based transgenes removed from their genomic context^18,21^ fail to assess effects on splicing or transcript stability, as well as potential artifacts of overexpression^26^. Genome editing technologies provide a means to overcome these challenges. Yet to our knowledge, genome editing has not yet been applied to functionally characterize VUS in *BRCA1* or other genes similarly linked to cancer predisposition.

Here we set out to apply genome editing to measure the functional consequences of all possible SNVs in *BRCA1*, regardless of whether they have been previously observed in a human. Given *BRCA1*’s immense size, this initial study focuses on 13 exons that encode the functionally critical RING and BRCT domains. In each experiment, a single exon is subjected to ‘saturation genome editing’^27^, wherein all possible SNVs are simultaneously introduced to a haploid human cell line in which *BRCA1* is essential. Consequently, *BRCA1* variants that result in nonfunctional alleles are depleted over time, a selection that is quantified by deep targeted sequencing. We optimized this method to obtain function scores for 3,893 SNVs, comprising 96.5% of all possible SNVs in the targeted exons. These function scores are bimodally distributed and nearly perfectly concordant with expert-based assessments of pathogenicity. We predict that our functional classifications will be of immediate clinical utility, and argue that the scaling of this approach to additional clinically actionable genes will substantially enhance the utility of genetic testing.

## RESULTS

### Saturation genome editing of *BRCA1* exons

Many genes in the HDR pathway, including those associated with hereditary cancer predisposition such as *BRCA1*, *BRCA2*, *PALB2* and *BARD1*^12^, were recently identified in a gene trap screen as being essential in the human haploid cell line HAP1^28^ (**Fig. 1a**). To validate this finding, we designed guide RNAs (gRNAs) to target exons of each of these genes and assessed HAP1 cell viability after transfecting each gRNA on a plasmid co-expressing Cas9 and a puromycin resistance cassette^29^. High cell death was evident by light microscopy (**Fig. 1b**), and a luminescence-based survival assay established that targeting any of these genes substantially reduces viability of HAP1 cells within one week (**Extended Data Fig. 1**). Deep sequencing of the edited loci of *BRCA1*-targeted cells confirmed that cell death was consequent to mutations, as there was widespread selection against frameshifting indels in favor of unedited loci and some in-frame indels (**Fig. 1c**). Overall, these results confirm the essentiality of HDR pathway components in HAP1 cells and establish targeted sequencing as a strategy to distinguish functional vs. non-functional *BRCA1* variants in a population of edited HAP1 cells.

**Figure 1.**
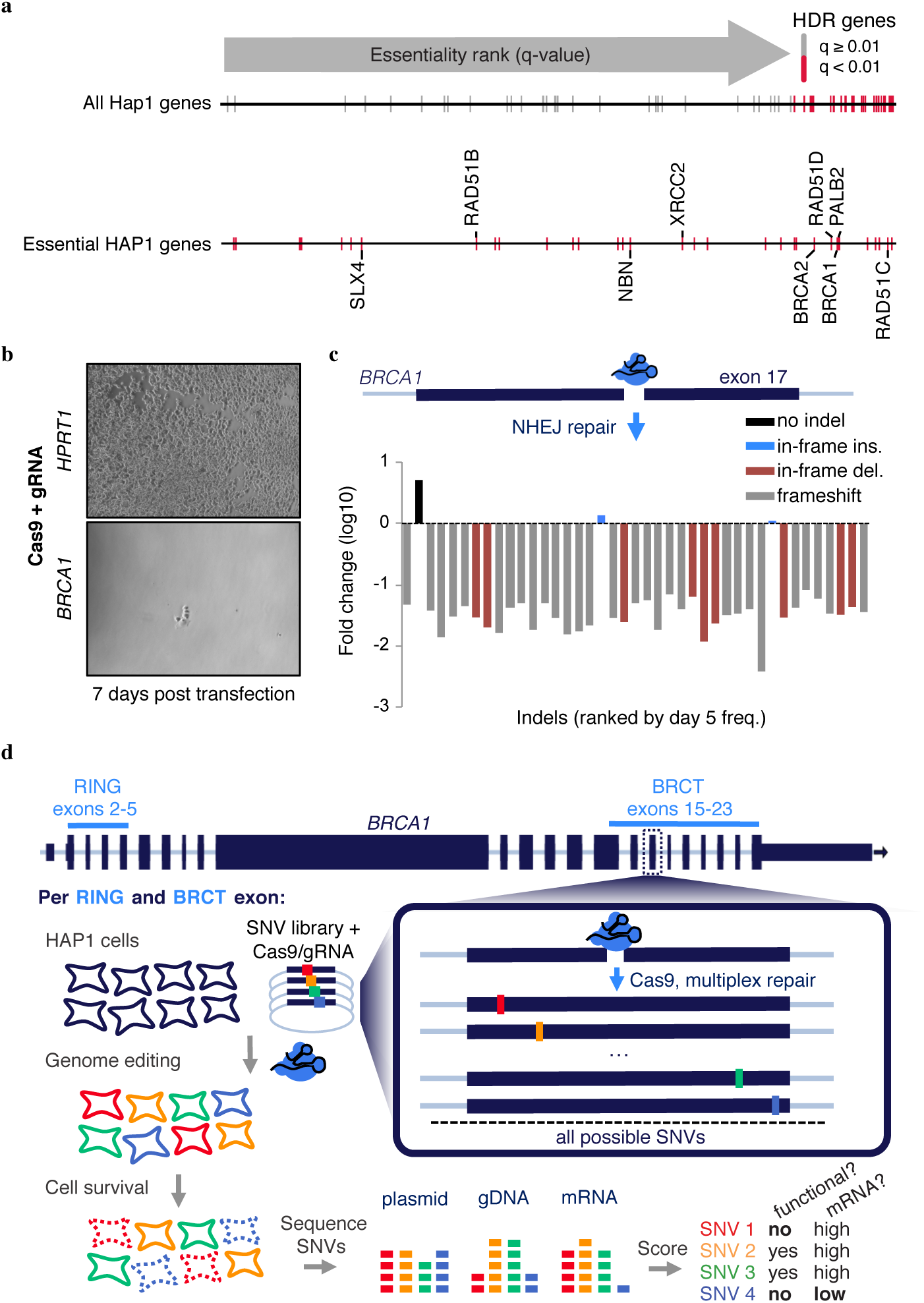
*BRCA1* and other HDR pathway genes are essential in HAP1 cells. **a**, The q-value rankings of HDR pathway genes (N = 66, defined by Gene Ontology) among 14,306 genes scored in a HAP1 gene trap screen for essentiality^28^ are indicated with tick marks. Essential HDR genes are colored red and those implicated in cancer predisposition are labelled in the enlargement below. Of the 66 HDR pathway genes scored, 34 including *BRCA1* were ‘essential’, a 3.4-fold enrichment compared to non-HDR genes (Fisher’s exact *P* = 6.1 × 10^−12^). **b**, HAP1 cell populations were transfected with a Cas9/gRNA plasmid either targeting the non-essential gene *HPRT1* (control) or exon 17 of *BRCA1* on day 0. Successfully transfected cells were selected with puromycin (days 1-4) and cultured until day 7, at which point cells were washed prior to imaging. Images are representative of two transfection replicates. **c**, The targeted *BRCA1* exon 17 locus was deeply sequenced from a population of transfected cells sampled on day 5 and day 11. The fold-change from day 5 to day 11 for each editing outcome observed at a frequency over 0.001 in day 5 sequencing reads is plotted. All alleles but indel-free sequences and two in-frame insertions were depleted. **d**, Saturation genome editing experiments were designed to introduce all possible SNVs across thirteen *BRCA1* exons encoding the protein’s RING (exons 2-5) and BRCT domains (exons 15-23). For each exon, a Cas9/gRNA construct was designed to be transfected with a library of plasmids containing all SNVs across ~100 bp of genomic sequence (the ‘SNV library’). SNV libraries were designed to saturate a total of 1,345 bp of genomic sequence, spanning BRCT and RING domain coding regions and adjacent intronic sequences. SNV library plasmids contain homology arms to mediate genomic integration, as well as fixed synonymous variants within the CRISPR target site to prevent Cas9 re-cutting. Upon HAP1 cell transfection of each Cas9/gRNA plasmid / SNV library pair, successfully edited cells harbor a single *BRCA1* SNV from the library. Cells are sampled 5 and 11 days after transfection and targeted gDNA and RNA sequencing is performed to quantify SNV abundances. SNVs compromising *BRCA1* function are selected against, manifesting in reduced gDNA representation, and SNVs impacting mRNA production are depleted in RNA samples relative to gDNA.

We next designed and optimized experiments for saturation genome editing (SGE)^27^ (**Fig. 1d**). We chose to focus on the thirteen exons of *BRCA1* encoding the RING (exons 2-5) and BRCT domains (exons 15-23) because these domains are essential for the protein’s role as a tumor suppressor^30–32^ and harbor missense variants known to be pathogenic or benign, as well as ~400 VUS or variants with conflicting reports of pathogenicity^13,33,34^. To create a library of repair templates, we used array-synthesized oligo pools containing all possible SNVs spanning each exon and ~10 bp of adjacent intronic sequence. Oligo pools for each exon were PCR-amplified and cloned into plasmids with homology arms to mediate genomic integration and make ‘SNV libraries’. Each SNV library molecule also included a fixed synonymous substitution at the target site to reduce re-cutting by Cas9 after successful HDR^27^. Each SGE experiment targeted a single exon. In brief, a population of 20 million HAP1 cells was co-transfected on day 0 with the exon’s corresponding SNV library and Cas9/gRNA plasmid. Successfully transfected cells were selected with puromycin (days 1-4), expanded, and sampled on day 5 and day 11. Variant frequencies were quantified by targeted amplification and sequencing of the edited exon from genomic DNA (gDNA) harvested on day 5 and day 11. Negative controls were used to confirm that PCR amplicons were not derived from the plasmid DNA of the SNV library.

We initially performed SGE experiments in replicate for each exon in wild-type (WT) HAP1 cells. In each of the 13 exons, we observed depletion of frameshifting indels, confirming intolerance to loss of *BRCA1* function (**Extended Data Fig. 2**). However, towards achieving more robust data, we optimized SGE in HAP1 cells in two ways. First, to increase HDR rates in HAP1 cells, we generated a monoclonal *LIG4* knockout HAP1 line (HAP1-Lig4KO) (**Extended Data Fig. 3a-b**). LIG4 acts in the non-homologous end joining (NHEJ) pathway, and its depletion can increase the proportion of cells with HDR-mediated repair of double-stranded breaks^35,36^. We observed a median 3.6-fold increase in HDR rates on day 5 in HAP1-Lig4KO relative to WT HAP1 (**Fig. 2a**). Second, it is known that HAP1 cells can spontaneously revert to diploidy^37^. Simply sorting HAP1 cells for 1N ploidy prior to editing improved reproducibility (**Extended Data Fig. 3c-e**).

**Figure 2.**
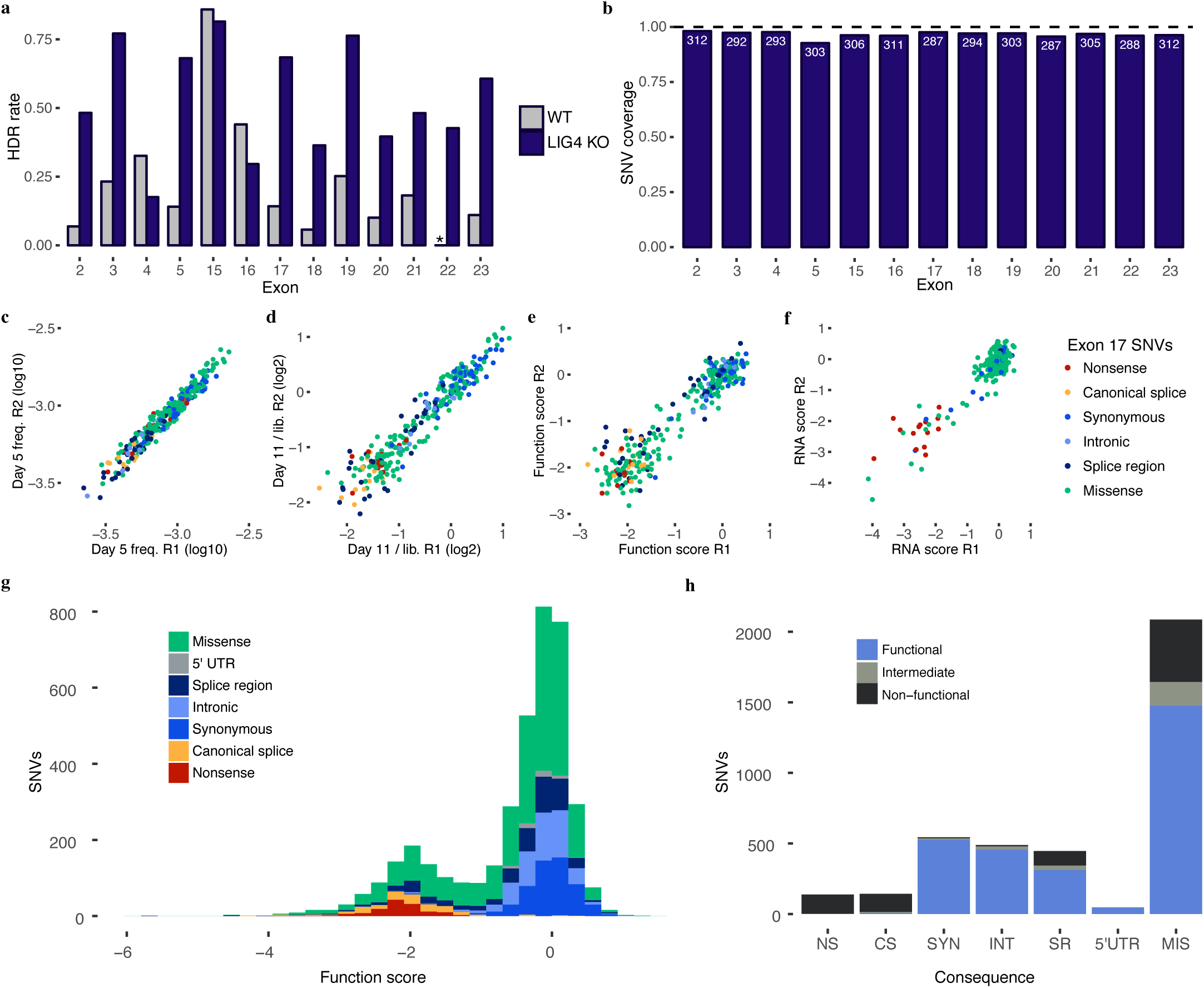
Saturation genome editing enables functional classification of 3,893 *BRCA1* SNVs. **a**, HDR editing rates were calculated for each exon as the fraction of day 5 reads containing the SNV library’s fixed synonymous variant (*i.e.* an ‘HDR marker’ edit). The average of two WT HAP1 replicates and two HAP1-Lig4KO replicates is plotted for comparison. (Asterisk denotes missing exon 22 data.) **b**, The fraction of all possible SNVs scored is shown for each exon. SNVs were excluded mainly due to proximity to the HDR marker and/or poor sampling (Extended Data Fig. 6 and Methods). **c-f**, Reproducibility was assessed across all exon replicates (Extended Data Fig. 5). Measurements for exon 17 SNVs assayed in HAP1-Lig4KO cells are plotted to show correlations of day 5 frequencies (**c**, ρ = 0.97), day 11 over library ratios (**d**, p = 0.95), function scores (**e**, p = 0.88), and RNA expression scores (**f**, ρ = 0.61). **g**, A histogram of 3,893 SNV function scores (averaged across replicates and normalized across exons) shows how each category of mutation compares to the overall distribution. **h**, The number of SNVs within each category of mutation is plotted and colored by functional classification determined by SGE. (NS = nonsense, CS = canonical splice, SYN = synonymous, INT = intronic, SR = splice region, 5’UTR = 5’ untranslated region, MIS = missense.)

We next performed optimized SGE experiments for each of the 13 targeted exons in 1N-sorted HAP1-Lig4KO cells, testing nearly every possible SNV per exon in replicate (**Fig. 2b**). Functional effects of SNVs on survival were determined by targeted DNA sequencing of each SNV library as well as the edited exon in gDNA harvested on day 5 and day 11 (**Fig. 2c-e**). Additionally, targeted RNA sequencing of day 5 samples was used to determine how abundant exonic SNVs were in *BRCA1* mRNA (**Fig. 2f**). Because these optimizations resulted in greater reproducibility (**Extended Data Fig. 4**), we moved forward with data from the 1N-sorted HAP1-Lig4KO cells only.

## Function scores for 3,893 *BRCA1* SNVs

We sought to calculate function scores for each SNV in a way that accurately quantified selection throughout the experiment while also minimizing experimental biases. First, we calculated the log2 ratio of the SNV’s frequency on day 11 vs. its frequency in the original plasmid library. Second, positional biases in editing rates were modeled (using day 5 SNV frequencies) and subtracted (**Extended Data Fig. 5**). Third, to enable comparisons between exons, we normalized function scores such that each experiment’s median synonymous and nonsense SNV matched global medians. Finally, a small number of SNVs were filtered out that could not confidently be scored (*e.g.* SNVs poorly represented on day 5; **Extended Data Fig. 6**). Altogether, we obtained function scores for 3,893 SNVs within or immediately intronic to these exons (**Fig. 2e, Supplementary Table 1**, https://sge.gs.washington.edu/BRCA1). This corresponds to 96.5% of all possible SNVs in these regions.

Function scores for SNVs in these 13 *BRCA1* exons were bimodally distributed (**Fig. 2g**). All nonsense SNVs scored below –1.25 (N = 138, median = −2.12), whereas 98.7% of synonymous SNVs >3 bp from splice junctions scored above −1.25 (N = 544, median = 0.00). We classified all SNVs as ‘functional’, ‘non-functional’, or ‘intermediate’ by fitting a two-component Gaussian mixture model in which the parameters of the ‘non-functional’ distribution were based on all nonsense SNVs and the ‘functional’ distribution based on synonymous SNVs not depleted in RNA (**Extended Data Fig. 7**). We then used this model to estimate the probability of each SNV’s score being drawn from the non-functional distribution (*P*_nf_). SNVs with *P*_nf_ < 0.01 were categorized as functional (72.5%); SNVs with *P*_nf_ > 0.99 were categorized as non-functional (21.1%); and SNVs with 0.01 < *P*_nf_ < 0.99 (6.4%) were categorized as intermediate.

Rare missense variants in *BRCA1* are particularly challenging to interpret clinically. Of the missense SNVs that we scored here, 21.1% (441 of 2,086) scored as non-functional (**Fig. 2h**). Although most of the remaining missense SNVs were functional (70.6%), there was an enrichment for missense SNVs with intermediate effects (8.1%, compared to 4.4% of all other SNVs; Fisher’s exact *P* = 2.7 × 10^−6^).

An advantage of assaying variants by genome editing is that their impact on native regulatory mechanisms such as RNA splicing can be ascertained^27^. Whereas SNVs disrupting canonical splice sites (the two intronic positions immediately flanking each exon) overwhelmingly scored as non-functional (89.5%) or intermediate (5.5%) (‘CS’ in **Fig. 2h**). SNVs positioned 1-3 bp into the exon or 3-8 bp into the intron had variable effects. We defined SNVs in these regions that did not alter the amino acid sequence as ‘splice region’ variants, of which 22.9% were non-functional (‘SR’ in **Fig. 2h**), on par with missense SNVs (21.2% non-functional). SNVs positioned more deeply in introns or in the 5’ UTR were similar to non-splice-region synonymous SNVs, in that they were much less likely to score as non-functional (intronic: 1.8% non-functional; 5’ UTR: 0.0% non-functional; synonymous: 1.3% non-functional).

## Function scores are nearly perfectly concordant with ClinVar

We next asked how well our function scores agreed with expert-based clinical variant interpretations, where available in ClinVar. Of 169 SNVs deemed ‘pathogenic’ in ClinVar that overlapped with our classifications, 162 were designated ‘non-functional’, 2 ‘functional’, and the remaining 5 ‘intermediate’. In contrast, of 22 SNVs deemed ‘benign’ in ClinVar that overlapped with our classifications, 1 was designated ‘non-functional’, 1 ‘intermediate’, and 20 ‘functional’ (**Fig. 3a**). The three SNVs for which our function scores are unambiguously discordant with ClinVar are discussed further below. A ROC curve showed a sensitivity of 96.7% at 98.2% specificity when we treat ‘likely pathogenic’ and ‘likely benign’ ClinVar annotations as pathogenic and benign, respectively (**Fig. 3b**). Importantly, our assay accurately predicts ClinVar interpretations independent of mutational consequence; sensitivity and specificity are high for both missense and splice site SNVs when these are considered separately from nonsense SNVs (**Extended Data Fig. 7f**). We find 64 of 256 (25.0%) VUS and 60 of 122 (49.2%) SNVs with conflicting interpretations to be non-functional in our assay (**Fig. 3c**). Missense VUS from ClinVar were significantly more likely to score as non-functional compared to missense SNVs absent from ClinVar (25.9% vs. 17.2%, *P* = 0.002). Apart from largely corroborating established ClinVar annotations, our scores also provide functional classifications for an additional 3,140 SNVs, the vast majority of which have yet to be publicly reported in clinical sequencing. Of these SNVs, 498 (15.9%) are classified as non-functional.

**Figure 3.**
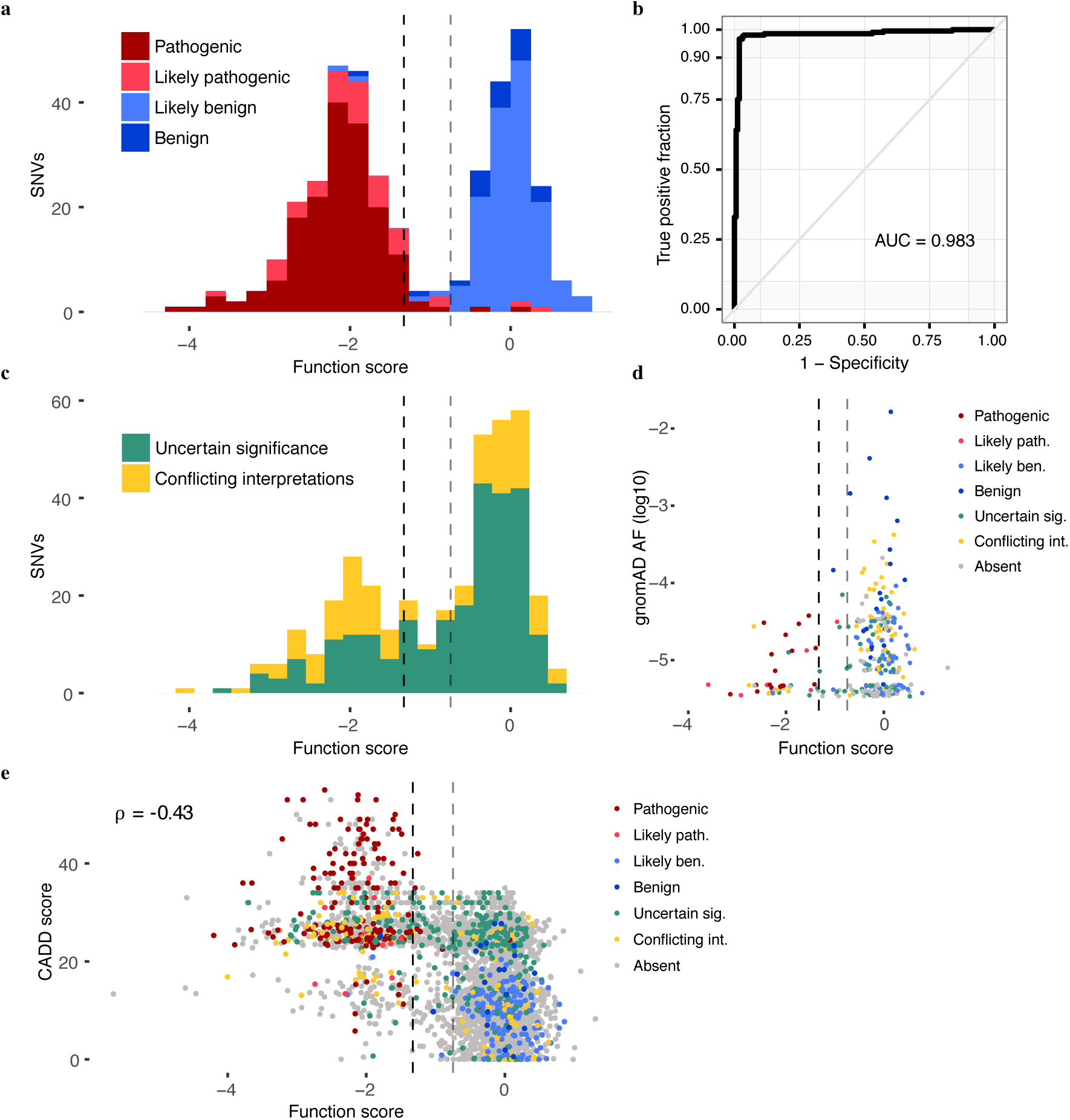
SGE function scores are highly accurate at predicting clinical interpretations of *BRCA1* SNVs. **a**, The distribution of SNV function scores colored by ClinVar interpretation. Scores are shown for the 375 SNVs with at least a ‘1-star’ review status in ClinVar and either a ‘pathogenic’ or ‘benign’ interpretation (including ‘likely’). The dashed lines indicate the functional classification thresholds determined by mixture modeling (gray = intermediate, black = non-functional). **b**, An ROC curve reveals optimal sensitivity and specificity for classifying the same 375 SNVs in **a** at SGE function score cutoffs from −1.03 to −1.22. **c**, The distribution of scores plotted as in **a** for the 378 SNVs annotated as variants of uncertain significance or with conflicting interpretations. 91.3% of such variants are classified as ‘functional’ or ‘non-functional’ by SGE. **d**,**e**, SNVs are colored by ClinVar annotation. **d**, Among the 302 SNVs assayed also present in gnomAD, higher allele frequencies associated with higher function scores (Wilcoxon Signed Rank Test, *P* = 3.7 × 10^−12^). **e**, CADD scores (which predict deleteriousness) inversely correlate with function scores.

We also investigated the relationship between our function scores and SNV frequencies in large-scale databases of human genetic variation. Of 302 assayed SNVs that overlap with the Genome Aggregation Database (gnomAD)^16^, higher allele frequencies were associated with higher function scores (**Fig. 3d**). For instance, 33 of 166 (19.9%) of singleton gnomAD variants were non-functional, whereas only 8 of 136 SNVs (5.9%) seen in multiple individuals were non-functional (Fisher’s exact *P* = 3 × 10^−4^). A similar trend was observed with the Bravo database (**Extended Data Fig. 8a**). The FLOSSIES database contains *BRCA1* variants observed in women over seventy years old who have not developed breast or ovarian cancer^38^. Of 39 intersecting SNVs, only one scored as non-functional (**Extended Data Fig. 8b**). Collectively, these observations show that *BRCA1* SNVs with higher allele frequencies are more likely to be functional, as expected. However, the fact that >70% of ClinVar variants and >95% of non-ClinVar variants that we assayed here have not been observed even once in sequencing of >120,000 humans illustrates the challenges facing observational approaches to variant interpretation.

Several computational metrics are currently used to the assess deleteriousness of variants and often included in genetic testing reports. Although our function scores correlate with metrics such as CADD^39^, phyloP^40^, and Align-GVGD^41^, which are largely based on evolutionary conservation and biochemical properties of missense variants, the modesty of these correlations underscores the value of functional assays (**Fig. 3e, Extended Data Fig. 9a-g**). ROC curve analysis restricted to missense variants reveals that SGE-based function scores outperform these metrics at predicting pathogenicity status in ClinVar (**Extended Data Fig. 9h-l**). This outperformance is likely underestimated because some of these metrics (*e.g.* Align-GVGD) or their correlates (*e.g.* evolutionary conservation) informed the ClinVar classifications of pathogenicity in the first place.

### Mechanisms of *BRCA1* loss-of-function

To gain insights into the various mechanisms by which SNVs compromise function, we performed targeted RNA sequencing of *BRCA1* transcripts from day 5 cells. We normalized SNV frequencies in cDNA to their frequency in gDNA to produce mRNA expression scores (‘RNA scores’) for 96% of the functionally characterized exonic SNVs. Together with function scores, RNA scores enable fine mapping of molecular consequences of SNVs (**Fig. 4**). For instance, regions of exons 2 and 15 that respectively code for RING and BRCT domain residues contain numerous loss-of-function missense variants. This contrasts with coding sequence in the same exons that fall outside of the boundaries of these protein domains. Overall, 89% of non-functional missense SNVs did not reduce RNA levels substantially, suggesting that their effects are likely mediated at the protein level (**Fig. 5a**). Many residues that are sensitive to missense SNVs *not* impacting RNA levels map to buried hydrophobic residues or to the zinc-coordinating loops that are required for proper RING domain folding (**Fig. 5b-c**). However, 11% of non-functional missense SNVs are depleted from RNA by 4-fold or more. Many of these SNVs map outside of key protein-protein interfaces and rather in unstructured loops, suggesting that they cause loss-of-function by lowering mRNA expression levels. Consistent with this, the 12 synonymous SNVs classified as non-functional also tended to markedly reduce mRNA levels (median 5.4-fold reduction).

**Figure 4.**
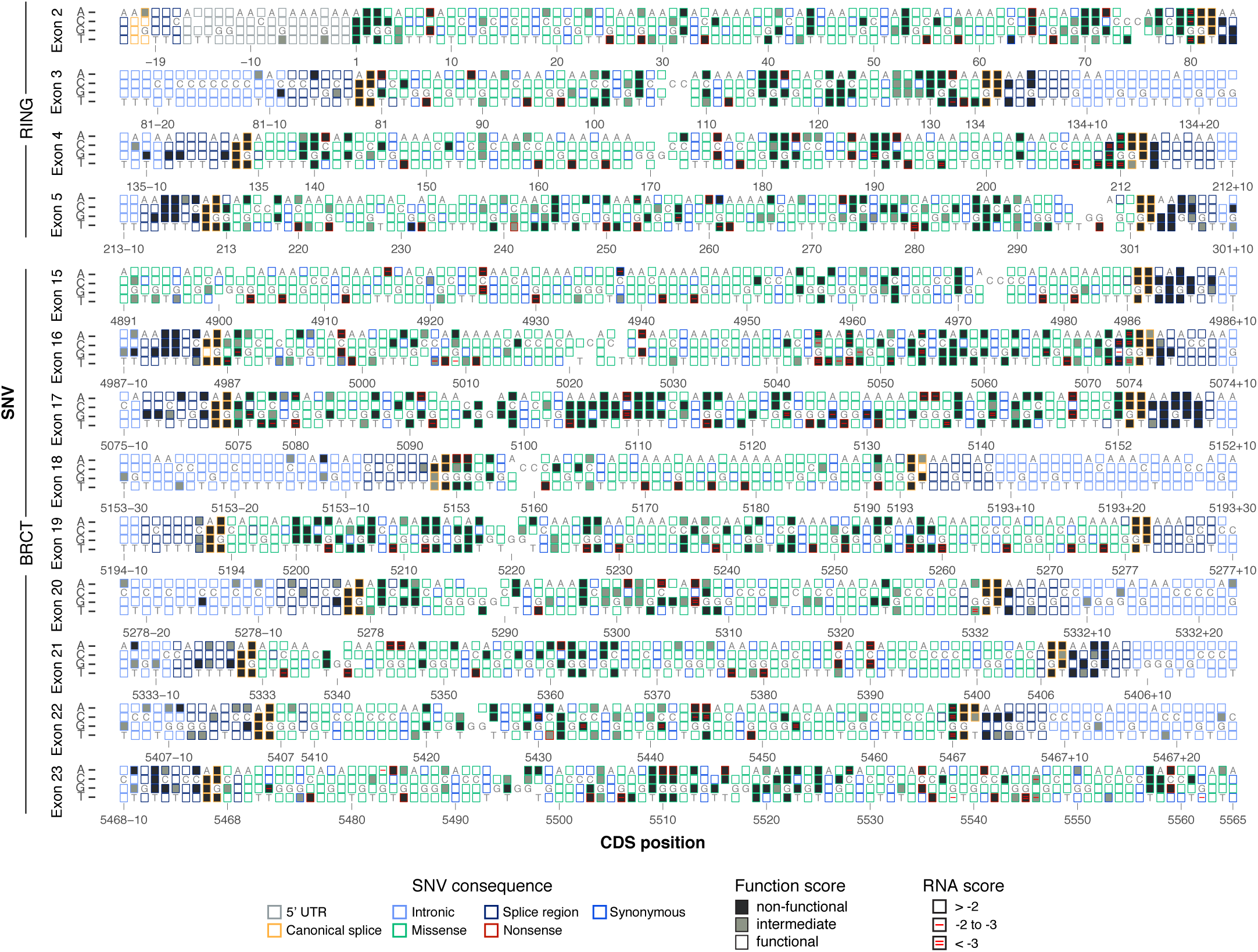
Sequence-function maps for 13 *BRCA1* exons. The 3,893 SNVs scored with SGE are each represented by a box corresponding to coding sequence position (NCBI transcript ID: NM_007294.3) and nucleotide identity. Boxes are filled corresponding to functional class, and outlined corresponding to the SNV’s mutational consequence. Red lines within boxes mark SNVs depleted in RNA; one line indicates an RNA score between −2 and −3 (log2 scale) and two lines indicate a score below −3. RNA measurements were determined only for exonic SNVs, excluding exon 18. Reference nucleotides are indicated by dark gray letters; blank boxes indicate missing data.

**Figure 5.**
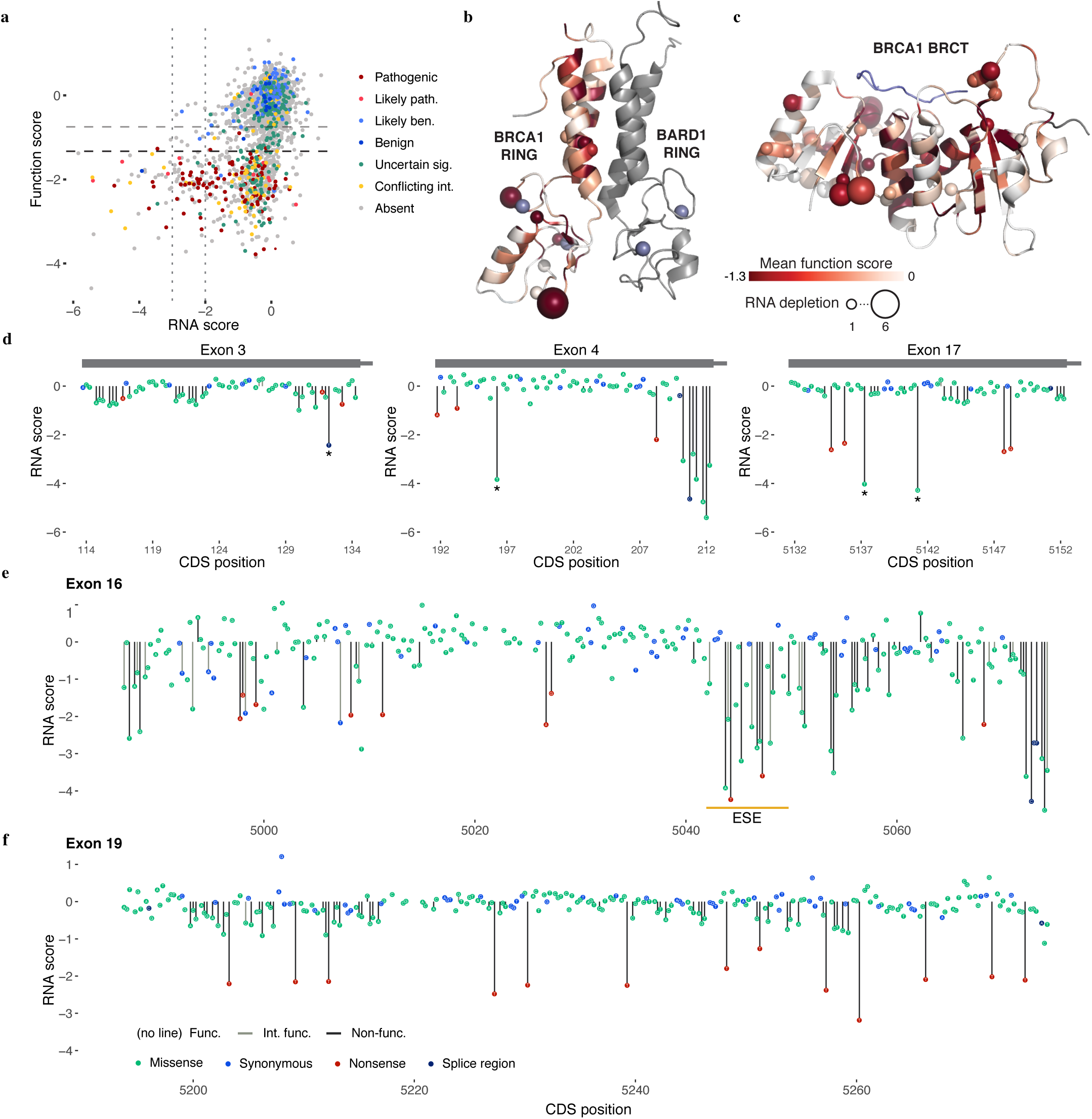
Measuring SNV mRNA abundance and function in parallel delineates mechanisms of variant effect. **a**, Function scores are plotted against RNA scores for all exonic synonymous and missense SNVs scored (N = 2,646). Horizontal dashed lines indicate functional thresholds, and vertical dotted lines mark RNA scores of −2 and-3. **b,c**, Function scores for all SNVs were mapped onto the structures of the RING (**b**, pdb 1JM7) and BRCT (**c**, pdb 1T29) domains in shades of red by averaging missense SNV scores at each amino acid position. The number of SNVs that cause >75% reduction in RNA levels at each amino acid position is represented by the size of the sphere at the alpha-carbon at each residue. Grey denotes residues not assayed and the BACH1 peptide bound to the BRCT structure is colored slate blue. **d**,**e**,**f**, SNV RNA scores are plotted by transcript position, with lines denoting SNV functional classification. **d**, Examples of non-functional SNVs with low RNA scores that create new 5’-GU splice donor motifs are shown. Complete maps of RNA scores for exons 16 (**e**) and exon 19 (**f**) reveal highly variable sensitivity to RNA depletion. The location of the strongest predicted exonic splice enhancer in exon 16^42^ is indicated by the orange line (**e**).

How do these exonic SNVs cause reductions in mRNA levels? Although other mechanisms cannot be ruled out, many of the variants depleted in mRNA are likely impacting RNA splicing. This is evidenced by an overrepresentation of non-functional SNVs near splice junctions, including low scores for many SNVs at terminal G nucleotides of exons (**Fig. 4**), non-functional exonic SNVs with low mRNA levels that create new acceptor or donor sequences (SNVs annotated with asterisks in **Fig. 5d**), and the presence of short regions (~6-8 bp) in which many SNVs have moderate-to-strong effects on RNA levels, suggestive of exonic splice enhancers^42^ (**Fig. 5e**). Certain exons appeared particularly prone to harbor non-functional SNVs with low RNA scores. In exon 16, for instance, 46 of 244 SNVs (excluding nonsense) were non-functional (**Fig. 5e**). Of these, more than half (n = 26) reduced RNA levels by more than 2-fold, and nearly a third (n = 15) by more than 4-fold. In contrast, in exon 19, of 55 of 234 SNVs (excluding nonsense) that were non-functional, none lowered expression by more than 2-fold (**Fig. 5f**). Exon 19 also completely lacks non-functional SNVs in its flanking intronic regions (apart from the acceptor and donor sites), suggesting the exon is robustly spliced compared to other exons.

### Discordances with ClinVar Interpretations

We leveraged sequence-function maps in reviewing the evidence around the three SNVs for which our classifications were clearly discordant with ClinVar. Discordant SNVs assayed in our preliminary experiments in WT HAP1 cells had similar scores, suggesting their classifications are not secondary to noise in our assay (**Extended Data Fig. 10**). One missense SNV designated ‘pathogenic’ in ClinVar that we scored as functional, c.5359T>A (C1787S), was identified through segregation with disease. However, in each case, it was seen in *cis* with a second SNV at the neighboring amino acid position^43^. Our data as well as data from other functional assays^44^ suggest c.5359T>A on its own is functional. The linked SNV c.5363G>T (G1788D), however, scored as non-functional, calling into question the ClinVar annotation (**Extended Data Fig. 10c**).

A second disagreement was identified in the exon 2 splice acceptor, c.-19-2A>G. This SNV was annotated as ‘pathogenic’ in ClinVar based on its occurrence at a splice acceptor site^45^, rather than from having been associated with disease. Exon 2 contains the *BRCA1* translation initiation codon, meaning that alternate splice forms may preserve the complete open reading frame. Of note, CADD scores for SNVs across the exon 2 acceptor site were much lower than for SNVs in other canonical splice sites (**Extended Data Fig. 10d**), and none of the 6 SNVs that we introduced here scored as non-functional. Further supporting that this splice site is not essential for *BRCA1* function, RNA sequencing from breast and ovarian tissue in the GTEx database^46^ shows this exon junction is poorly represented among *BRCA1* transcripts (**Extended Data Fig. 10e**). This suggests that this acceptor site is likely dispensable both in our assay and in tissues relevant to disease, again calling the ClinVar annotation into question.

Exon 16 harbored the third discordantly classified SNV, the ‘benign’ c.5044G>A (E1682K) variant, which scored as non-functional in our assay. Of note, c.5044G>A resides in a predicted exonic splice enhancer (ESE)^42^, and its low function score was substantiated by a reduction in RNA levels of over 90% (**Fig. 5e**). Neighboring SNVs in the predicted ESE also reduced RNA expression, corroborating the element’s importance. Although this missense SNV is rare (absent from gnomAD and Bravo), reports indicate it was designated as benign based on being observed in *trans* with a variant considered pathogenic^33^, as biallelic *BRCA1* loss-of-function mutations are thought to be embryonic lethal. The underlying data supporting this finding are not publically available, and previous assays of this variant did not measure splicing consequences^44^.

## DISCUSSION

Here we applied saturation genome editing to the 13 exons that encode functionally critical domains of the cancer risk gene, *BRCA1*, characterizing the functional consequences of nearly 4,000 SNVs in their native genomic context. Specifically, we used CRISPR/Cas9 to introduce hundreds of SNVs per experiment, followed by deep sequencing to measure the functional consequences of each SNV in parallel. Because we measured cell survival, the effects of SNVs on multiple layers of gene function (*e.g.* RNA splicing, translation, protein function, protein stability) are effectively integrated. The approach is validated by nearly perfect concordance of function scores with available evidence for clinical pathogenicity.

Our experimental approach has several caveats. First, the exact requirements for *BRCA1* function essential to maintaining *in vitro* viability and growth of HAP1 cells, as opposed to mediating *in vivo* tumor suppression, are not known. For instance, we cannot rule out, differences in splicing or dosage requirements between our *in vitro* model vs. *in vivo* physiology. Second, we are not currently able to interrogate every possible SNV. Of note, most of the 3.5% of SNVs for which we do not provide function scores were excluded by factors related to genome editing, rather than because of sampling (**Extended Data Fig. 6**). Lastly, as these experiments were designed to measure loss-of-function in a haploid cell line, we are unable to detect all types of functional effects (*e.g.* dominant negative variants).

Notwithstanding these limitations, we achieved nearly comprehensive coverage of the targeted regions and our functional classifications are nearly perfectly concordant with current clinical interpretations. As such, we anticipate that our results will be clinically useful, both for adjudicating hundreds of observed variants whose interpretation is currently ambiguous, as well as for providing immediate functional assessments for variants newly observed. Therefore, the pressing question becomes how to best to integrate this functional data within existing clinical variant classification schemes^47^.

A benefit of functional data is that measurements are systematically derived, independent of prior expectation^48^. As such, function scores add an additional layer of evidence to support interpretations of variants made through segregation with disease. However, for the large number of VUS for which genetic evidence is insufficient, the predictive power demonstrated here suggests function scores can be used to classify variants with >95% accuracy. As current standards for defining ‘likely pathogenic’ and ‘likely benign’ variants accept a comparable level of uncertainty^49^, we argue that a failure to use appropriately validated functional data to inform clinical care would be a missed opportunity. There is precedent for incorporating functional data in interpretation guidelines^24^, but the breadth and predictive power demonstrated by SGE calls for an increased role. Indeed, given the low likelihood that observational approaches will ever be sufficient to classify variants not yet seen once in humans, we believe that there is a strong argument to be made for using highly predictive function scores, where available, to inform initial interpretations of newly observed variants.

The orthologous nature of SGE data also presents an opportunity for integration with other data sources. For example, a multiplex reporter assay for HDR activity strengthens the functional evidence presented here for *BRCA1* missense variants (see accompanying manuscript from Starita *et al*.). Integration and optimal weighting of experimental and computational approaches may also further improve classification of variants lacking genetic evidence. In cases where evidence is contradictory, functional data may yield specific hypotheses to test. For example, c.5044G>A, for which our data contradicts the ClinVar interpretation (**Fig. 5e**), would be disambiguated by testing *BRCA1* mRNA levels in individuals harboring this SNV. Similar approaches should be taken to more confidently resolve unlikely functional classifications, such as synonymous SNVs with low function scores and canonical splice SNVs deemed functional. Furthermore, the ~6% of SNVs exhibiting intermediate function scores remain beyond definitive interpretation. The fact that we observe an excess of missense SNVs with intermediate scores suggests that some of these may be hypomorphic *BRCA1* alleles^50-52^. Further studies will be necessary to quantify the penetrance of intermediately functional variants.

Moving forward, our study provides a blueprint for comprehensive functional analysis of all potential SNVs in clinically actionable genes for which appropriate assays can be developed. Here, we prioritized *BRCA1* exons encoding the RING and BRCT domains, but SGE of the entire coding sequence and promoter are also well motivated. Furthermore, the essentiality of *BRCA2*, *PALB2*, *BARD1*, and *RAD51C* in HAP1 suggests that these genes are assayable by the same method. For genes in other pathways, assays that are compatible with saturation genome editing (*e.g.* drug selection, FACS on phenotypic markers, etc.) may need to be developed and validated. For any gene tested, it is critical that functional measurements be calibrated to clinical evidence of pathogenicity. Given that SGE tests variants in their endogenous genomic context, the scaling of SGE to many loci promises to improve our understanding of how diverse biological functions are encoded by the genome.

Delivering on the promise of genomic medicine requires that we not only be able to cost-effectively ascertain genetic variation, but also accurately and definitively interpret it. Presently, interpretation is the rate limiting step. As a potential path forward, we show that saturation genome editing is a viable strategy for functionally classifying thousands of variants in a clinically actionable gene, most of which have yet to be observed in a human. With further scaling, we anticipate that this paradigm will substantially improve the utility of genetic information in clinical decision making.

## DATA AVAILABILITY

Saturation genome editing data is available at: https://sge.gs.washington.edu/BRCA1.

## ACKNOWLEDGEMENTS

We thank Malte Spielmann, Daniela Witten, Aaron McKenna, Martin Kircher, Max Dougherty, John Lazar, Yi Yin, and Brian Shirts for insights on data analysis and/or comments on the manuscript, Jacob Kitzman for sharing reagents and protocols, Rocío Acuña-Hidalgo and Jennifer Milbank for experimental assistance and the Feng Zhang lab for sharing Cas9/gRNA plasmids. This work was supported by an NIH Director’s Pioneer Award (DP1HG007811 to J.S.) and a training award from the National Cancer Institute (F30CA213728 to GMF). JS is an Investigator of the Howard Hughes Medical Institute.

## METHODS

### HDR pathway essentiality analysis in HAP1 cells

HAP1 cells were derived from KBM7 cells (a near-haploid immortalized chronic myelogenous leukemia line) by introduction of induced pluripotent stem cell factors^56^. HAP1 gene essentiality scores were obtained^28^ and filtered on genes with greater than 20 mapped gene-trap insertions (N = 14,306). Of 78 HDR genes defined by the GO term ‘double-strand break repair via homologous recombination’ (GO:0000724), 66 were among the 14,306 genes included in analysis. To rank genes by essentiality, they were first ordered by q-value (low to high) and second by the proportion of gene-trap insertions in the sense orientation (low to high). HDR pathway genes implicated in cancer (labelled in Fig. 1) were defined as those included on the University of Washington BROCA sequencing panel^57^.

### gRNA design and cloning

All CRISPR gRNAs used in SGE and essentiality experiments were cloned into pX459^29^. This plasmid expresses the gRNA from a U6 promoter, as well as a Cas9-2A-puromycin resistance (puroR) cassette. *S. pyogenes* Cas9 target sites were chosen for SGE experiments on multiple criteria, assessed in the following order: 1.) To induce cleavage within *BRCA1* coding sequence, 2.) To target a genomic site permissive to synonymous substitution within the guanine dinucleotide of the PAM or the protospacer, 3.) To have minimal predicted off-target activity^58^, 4.) To have maximal predicted on-target activity^59^.

Complementary oligos ordered from Integrated DNA Technologies (IDT) were annealed, phosphorylated, diluted and ligated into BbsI-digested and gel-purified pX459, as described^29^. Ligation reactions were transformed into *E. coli* (Stellar competent cells, Takara), which were plated on ampicillin. Colonies were cultured and Sanger sequenced to confirm correct gRNA sequences. Purification of sequence-verified plasmids for transfection was performed with the ZymoPure Maxiprep kit (ZymoResearch). For targeting *LIG4* in HAP1 cells, pX458^29^ was used instead of pX459, which expresses EGFP in lieu of puroR.

### HDR library design and cloning

Array-synthesized oligos were designed as follows for each saturation genome editing region (*i.e.* a *BRCA1* exon). The sequence to be mutated (~100bp) was obtained from the human genome (hg19) and a synonymous substitution was introduced at the chosen Cas9 target site (*e.g.* a substitution at the PAM site). This ‘fixed’ substitution in the library was included in design to serve multiple purposes: 1.) plasmid library molecules harboring the substitution are predicted to be cleaved less frequently by Cas9:gRNA complexes, 2.) SNVs introduced to cells are predicted to be depleted via Cas9 re-cutting less frequently as a consequence of the fixed substitution, and 3.) sequencing reads can be filtered on the fixed substitution to distinguish true SNVs introduced via HDR from sequencing errors. A second synonymous substitution at an alternative CRISPR target site was introduced to the sequence as well, such that each exon’s SNV library would be compatible with multiple gRNAs. Next, a sequence was created for every possible single nucleotide substitution on this template. For all sequences, adapters were added to both ends to enable PCR amplification from the oligo pool. For each SGE region, the total number of oligos designed was three times the length of the region, plus the oligo template without any SNV (*e.g.* for a 100 bp SGE region, 301 total oligos were designed).

Pooled oligos were synthesized (Agilent Technologies). Primers designed to amplify the subset of oligos corresponding to a single exon’s region were used to perform PCR with Kapa HiFi Hot-start Ready Mix (‘Kapa HiFi’, Kapa Biosystems). PCR products were purified with Ampure beads (Agencourt) to be used in subsequent library cloning reactions.

Homology arms were cloned into pUC19 by PCR-amplifying (Kapa HiFi) regions surrounding each targeted exon from HAP1 gDNA. Primers for these reactions were designed such that homology arms would be between 600 and 1,000 bp on both sides of the targeted region. Adapters homologous to pUC19 were added to primers to facilitate NEBuilder HiFi Assembly cloning (NEB) into a linearized pUC19 vector. Cloning reactions were transformed into Stellar competent cells and selected with ampicillin. Plasmid DNA was isolated from colonies (Qiagen MiniPrep kit) and sequence-verified.

To make the HDR library, homology arm plasmids were linearized via PCR using primers that conferred 15-20 bp of terminal overlap with the adapter sequences flanking each PCR-amplified oligo pool. This sequence overlap enabled cloning via the NEBuilder HiFi Assembly Cloning Kit (NEB). Cloning reactions were transformed into Stellar competent cells, and a small proportion (1%) of the transformation was plated on ampicillin-containing plates to assess efficiency. All remaining transformed cells were grown directly in 100 ml of media with ampicillin for 16-18 hours, and plasmid DNA from the culture was isolated (ZymoPure Maxiprep kit) to produce each final HDR library.

### HAP1 cell culture

Quality-controlled WT HAP1 cells were purchased (Haplogen/Horizon Discovery) and cultured in media comprising Iscove’s Modified Dulbecco’s Medium (IMDM) with L-glutamine and 25 mM HEPES (GIBCO) supplemented with 10% fetal bovine serum (Rocky Mountain Biologicals) and 1% penicillin-streptomycin (GIBCO). Cells were grown on plates at 37C with 5% CO2, and passaged prior to becoming confluent. For routine passaging, cells were washed once with 1x phosphate buffered saline (PBS, Gibco), trypsinized with 0.25% trypsin with EDTA (Gibco), resuspended in media, centrifuged for 5 min at 300 rcf, and then resuspended and plated.

A monoclonal *LIG4* knock-out HAP1 line (HAP1-Lig4KO) was generated by transfecting a plasmid expressing a Cas9-2A-GFP cassette and a gRNA targeting the human *LIG4* coding sequence (gRNA sequence: 5’-GCATAATGTCACTACAGATC) into WT HAP1 cells. Single GFP-expressing HAP1 cells were sorted into wells of a 96-well plate and cultured. After two weeks, gDNA was harvested and Sanger sequencing was performed to assess *LIG4* editing. A clone with a 4bp deletion was identified and expanded further for use in saturation genome editing experiments.

HAP1 cells can spontaneously revert to a diploid state in cell culture. Therefore, to sort a 1N-enriched population of cells prior to transfection, cells were stained for DNA content with Hoechst 34580 (BD Biosciences) at 5 ug/ml media for 1h at 37C. FACS was performed to isolate 1-2×10^6^ cells from the lowest intensity Hoechst peak, corresponding to 1N ploidy. These cells were expanded for seven days prior to transfection.

### Transfection of HAP1 cells

For all experiments, HAP1 cells were transfected using TurboFectin 8.0 (Origene) according to manufacturer’s protocol. A 2.5x volume of Turbofectin was added to the transfection mix for each ug of plasmid DNA in Opti-Mem (Life Technologies). For each SGE transfection, 10 million cells were passaged to a 10 cm dish. The next day (day 0), cells were co-transfected with 12 ug of the Cas9/gRNA plasmid (pX459) and 3 ug of the SGE library corresponding to a single exon. For negative control transfections, a pX459 vector targeting *HPRT1* was used instead. On day 1, cells were passaged into media supplemented with puromycin (1 ug/ml) to select for successfully transfected cells. On day 4, cells were washed twice and passaged to 6 cm plates in regular media.

Cell populations were sampled on day 5 and day 11 for all SGE experiments. On day 5, half of the cells were pelleted and frozen and the other half passaged. The cells were passaged on day 8 into 15 cm dishes and then harvested on day 11. Negative control transfections were harvested on day 5.

For the luminescence-based viability assay, HAP1 cells were plated at ~35-40% confluency in a 6-well dish (approximately 1.2 million cells per well per target) then transfected with 1.5 ug Cas9/gRNA plasmid targeting coding exons of HDR genes or controls the following day. 24 hours after transfection the cells were plated in time-point triplicates at 20,000 cells per well in 96-well clear bottom plates in media with and without puromycin. Cells without puromycin were assessed 4 hours after plating to establish baseline absorbance for each target. Cell survival was assessed at day 2, day 5, and day 7 post-transfection using the CellTiterGlow reagent (Promega, 1:10 dilution of suggested reagent). Luminescence at 135 nm absorbance was measured using a Synergy plate reader (Biotek Instruments).

### Nucleic acid sampling and sequencing library production

For obtaining WT HAP1 genomic DNA for cloning homology arms and for genotyping the HAP1-Lig4KO cell line, DNA was isolated using the DNeasy kit (Qiagen). For each SGE experiment, DNA and total RNA were purified using the AllPrep kit (Qiagen). DNA samples were quantified with the Qubit dsDNA Broad Range kit (Thermo Fisher) and RNA samples by UV spectrometry (Nanodrop). PCR primers for genomic DNA were designed such that one primer would anneal outside of the homology arm sequence, thereby selecting for amplicons derived from gDNA and not plasmid DNA. PCR conditions were optimized using gradient qPCR on WT HAP1 gDNA.

All gDNA harvested from the population of day 5 cells was sampled by performing many PCR reactions in parallel on a 96-well plate, using 250 ng of gDNA per 50 ul reaction such that all day 5 gDNA was used in PCR (Kapa HiFi). At least as many PCR reactions were performed for day 11 samples (which yielded more gDNA) to ensure adequate sampling. PCRs were performed for the minimal number of cycles needed to complete amplification, with cycling conditions as specified in the Kapa HiFi protocol. An additional PCR was performed using day 5 gDNA from negative control transfections for each exon.

After PCR, multiple wells of amplicons from the same sample were pooled and purified using Ampure beads. Next, a nested qPCR was performed using the first reaction as template to produce a smaller amplicon with custom sequencing adapters (‘PU1L’ and ‘PU1R’), which was likewise purified with Ampure beads. The SGE libraries were also PCR-amplified at this step, starting from 50 ng of plasmid DNA. Lastly, a final qPCR was performed using purified products from the second reaction as template to add dual sample indexes and flow cell adapters.

RNA was sampled from day 5 HAP1-Lig4KO cells (AllPrep, Qiagen). Reverse transcription followed by RNase H treatment was performed on all RNA harvested or a maximum of 5 ug per sample (Superscript IV Kit, Life Technologies). This reaction was primed with a gene-specific primer complementary to the 3’ UTR in exon 23 of *BRCA1*. Primers were designed for each exon to amplify across exon junctions, and reaction conditions were optimized using gradient PCR. cDNA was distributed into 5 equal PCR reactions, which were run on a qPCR machine and then pooled in equal ratios. Flow cell adapters and sample indexes were added in an additional reaction (as for gDNA samples).

All sequencing libraries were purified with Ampure beads, quantified with the Qubit dsDNA High Sensitivity kit (Life Technologies), diluted and denatured for sequencing in accordance with protocols for the Illumina NextSeq or MiSeq machines.

### Sequencing and data analysis

Sequencing was performed on an Illumina NextSeq or MiSeq instrument, allocating about 3 million reads to each gDNA and cDNA sample, 1 million reads for each HDR library, and 500,000 reads for each negative control sample. gDNA samples for individual exons were sequenced on the same run. 300 cycle kits were used, with 150 cycles for read 1 and read 2 each, and 19 cycles for dual index reads. Custom sequencing primers and indexing primers are provided in Supplementary Table 2. Illumina PhiX control DNA was added to each sequencing run (~10% MiSeq, ~30-40% NextSeq) to improve base calling.

Illumina’s bcl2fastq 2.16 was used to call bases and perform sample demultiplexing and fastqc 0.11.3 was run on all samples to assess sequencing quality. SeqPrep was used with the following parameters to perform adapter trimming and to merge perfectly matched overlapping read pairs: ‘-A GGTTTGGAGCGAGATTGATAAAGT -B CTGAGCTCTCTCACAGCCATTTAG -M 0.1 -m 0.001 -q 20 -o 20’. Merged reads containing ‘N’ bases were removed. Reads from cDNA samples were removed if they contained indels or did not perfectly match transcript sequence flanking each targeted exon. Remaining cDNA reads were processed to match genomic DNA amplicons by removing flanking exonic sequence and replacing it with the exon’s corresponding intronic sequence. All reads were then aligned to reference gDNA amplicons for each exon using the needleall command in the EMBOSS 6.4.0 package with the following parameters: ‘-gapopen 10 -gapextend 0.5 -aformat sam’. Reads not aligning to the reference amplicon (alignment score < 300) were removed from analysis. To analyze indels, unique cigar counts were quantified from day 5 and day 11 samples using a custom Python script. Reads were classified as HDR events for rate calculations if the programmed edit or edits to the PAM or protospacer (HDR marker edits) were observed in the alignment. Variants without identifiable markers of HDR were not used. Abundances of SNVs were quantified only from aligned reads that had no other mismatches or indels, with the exception of the HDR markers. SNV reads with only the cut-site proximal HDR marker were summed with reads that had both HDR markers to get total abundances for each SNV in each sample, to which a pseudocount of 1 was added to all variants present in either the library, day 5 or day 11 sample. Frequencies for each SNV were calculated as SNV reads over total reads. SNV measurements from WT HAP1 cells and HAP1-Lig4KO cells were processed separately at all steps.

### Modeling positional biases of library integration

Positional biases in editing rates were modeled for each SNV by using a LOESS regression to fit the log2 day 5 over library ratios as a function of chromosomal position. To avoid modeling biological effects instead of positional effects, the model was fit only on the subset of SNVs that were not substantially depleted between any two timepoints in the experiment (*i.e.* SNVs with day 5 over library ratios > 0.5 and day 11 over d5 ratios > 0.8.). The regression was performed for each exon replicate, using the ‘loess’ function in R with span = 0.15. Each model was extended flatly outward to include any positions not fit (a total of 22 nucleotides of sequence on the edges of the edited regions). We subtracted each SNV’s positional fit (*e.g.* the model’s output) from the SNV’s log2 day 11 over library ratio to get position-adjusted ratios for each SNV.

### Normalizing scores within and across exons

Position-adjusted log2 day 11 over library ratios were normalized first across exon replicates, and then across all exons assayed. Scores from within each replicate were linearly scaled such that the median synonymous and median nonsense SNVs within the replicate were set to the median synonymous and median nonsense SNV values averaged across replicate experiments. The ensuing SNV scores for each replicate were then normalized across exons in the same way by again using median synonymous and median nonsense SNVs.

### SNV functional class assignment

Function scores were averaged across replicates and a mixture model was used to estimate the probability that each SNV’s score was drawn from the non-functional distribution of scores. The non-functional distribution was defined as nonsense SNVs across all exons. The functional distribution was defined as exonic synonymous SNVs not within 3 bp of splice junctions and with RNA scores within 1 standard deviation of the median synonymous SNV. This definition does not fully guarantee that these SNVs have no functional consequence. The means and variances of the ‘non-functional’ and ‘functional’ groups were fixed and a model was fit using the normalmixEM function of the mixtools package in R, with starting component proportions set to 0.5. The posterior probabilities generated from the model were used as point estimates of the probability of drawing each SNVs score from the non-functional distribution (*P*_nf_). Functional classifications were made by setting thresholds for *P*_nf_ as follows: *P*_nf_ > 0.99 = ‘non-functional’, 0.01< *P*_nf_ < 0.99 = ‘intermediate’, *P*_nf_ <0.01 = ‘functional’.

Independent of mixture modelling, ROC curves were used to assess performance of SGE data and other metrics’ ability to predict assigned ClinVar classifications. These analyses were performed with the plotROC package in R, and Youden’s J-statistic was calculated (sensitivity plus specificity minus 1) to determine optimal values reported in text.

### Variant filtering

A small minority of SNVs that could not be accurately scored were removed from analysis. If a SNV was not present in the HDR library at a frequency over 1 in 10^4^, it was presumed to have been lost in oligo synthesis or cloning and was removed. Additionally, if a SNV was not observed with complete HDR markers at a frequency over over 1 in 10^5^ in day 5 genomic DNA samples from both replicate experiments, it was removed. SNVs introduced near the CRISPR recognition site have the potential to facilitate Cas9 recutting of the locus (*e.g.* by replacing the PAM edit or introducing an alternative PAM site). Because these SNVs are likely to score lower consequent to Cas9 editing biases and not their effects on gene function, SNVs were filtered that created increased potential for re-cutting as follows: When an HDR marker mutation used to disrupt editing occurred at position 2 of the PAM (*e.g.* ‘NGG’ to ‘NCG’), SNVs that replaced this marker with an alternate base were removed to prevent biases introduced by recutting non-canonical *S. pyogenes* Cas9 PAMs (*e.g.* ‘NAG’, ‘NTG’). Additionally, variants that created a new PAM 1 bp 3’ of the mutated PAM were excluded due to the potential for recutting (*e.g.* unedited <u>PAM</u>: 5’-<u>NGG</u>A, edited PAM with HDR marker: 5’-<u>NCG</u>A, filtered out **SNV** that creates *new PAM* +1bp 3’: 5’-<u>N*CG*</u>**G**). (Extended Data Fig. 6 describes recutting observed at alternative PAMs.) To prevent misinterpretation, we also removed SNVs that created amino acid changes specific to the context of the library’s fixed edits (*e.g.* if in the unedited background, the SNV causes an X to Y change, but with a fixed edit in the same codon, the SNV causes an X to Z change). We also applied this logic to remove SNVs that introduced splice donor sites only in the context of the edited PAM, and SNVs that create splice donor sites in the unedited context but not in the context of the edited PAM.

The RNA scores for exon 18 samples were neither well correlated across replicates nor with SNV abundances in genomic DNA, indicating likely bottlenecking in library preparation. Therefore, RNA data from exon 18 was excluded. WT HAP1 function scores from exon 22 were excluded because there was an unusually high correlation between SNV frequencies sampled from the plasmid library and from day 5 gDNA, suggesting plasmid contamination in gDNA sequencing. This problem was fixed by designing a new primer to prepare gDNA sequencing samples from HAP1-Lig4KO cells.

### External data sources

Variant annotations were downloaded from CADD^39^ version 1.3 (http://cadd.gs.washington.edu/download). This included the following scores: mammalian phyloP, Grantham deviation, SIFT, Polyphen-2, and CADD. Align-GVGD scores were obtained by running the Align-GVGD program on BRCA1 sequences conserved to sea urchin. ClinVar data were downloaded on 1/2/2018 for all germline SNVs with at least a 1-star annotation. SNVs annotated as ‘Benign/Likely benign’ were grouped with ‘Likely benign’ SNVs and SNVs classified ‘Pathogenic/Likely pathogenic’ were grouped with ‘Likely pathogenic’ SNVs. SNV allele frequencies were obtained from http://gnomad.broadinstitute.org/ on 12/26/2017 for gnomAD^16^, from https://bravo.sph.umich.edu/freeze5/hg38/ on 11/19/2017 for Bravo, and from https://whi.color.com/ on 10/9/2017 for FLOSSIES data. Transcript data was obtained from GTEx on 1/3/2018. Throughout this study, *BRCA1* exons, coding nucleotide positions, and amino acid positions are referenced by the ClinVar transcript annotation for *BRCA1*, transcript NM_007294.3 (NCBI).

### Statistical reporting

All statistical tests described were performed as two-tailed tests using the R software package.

### Code availability

Custom scripts for analyzing sequencing data were written in Python and R. All code will be made available upon request.

---

**Extended Data Figure 1.**
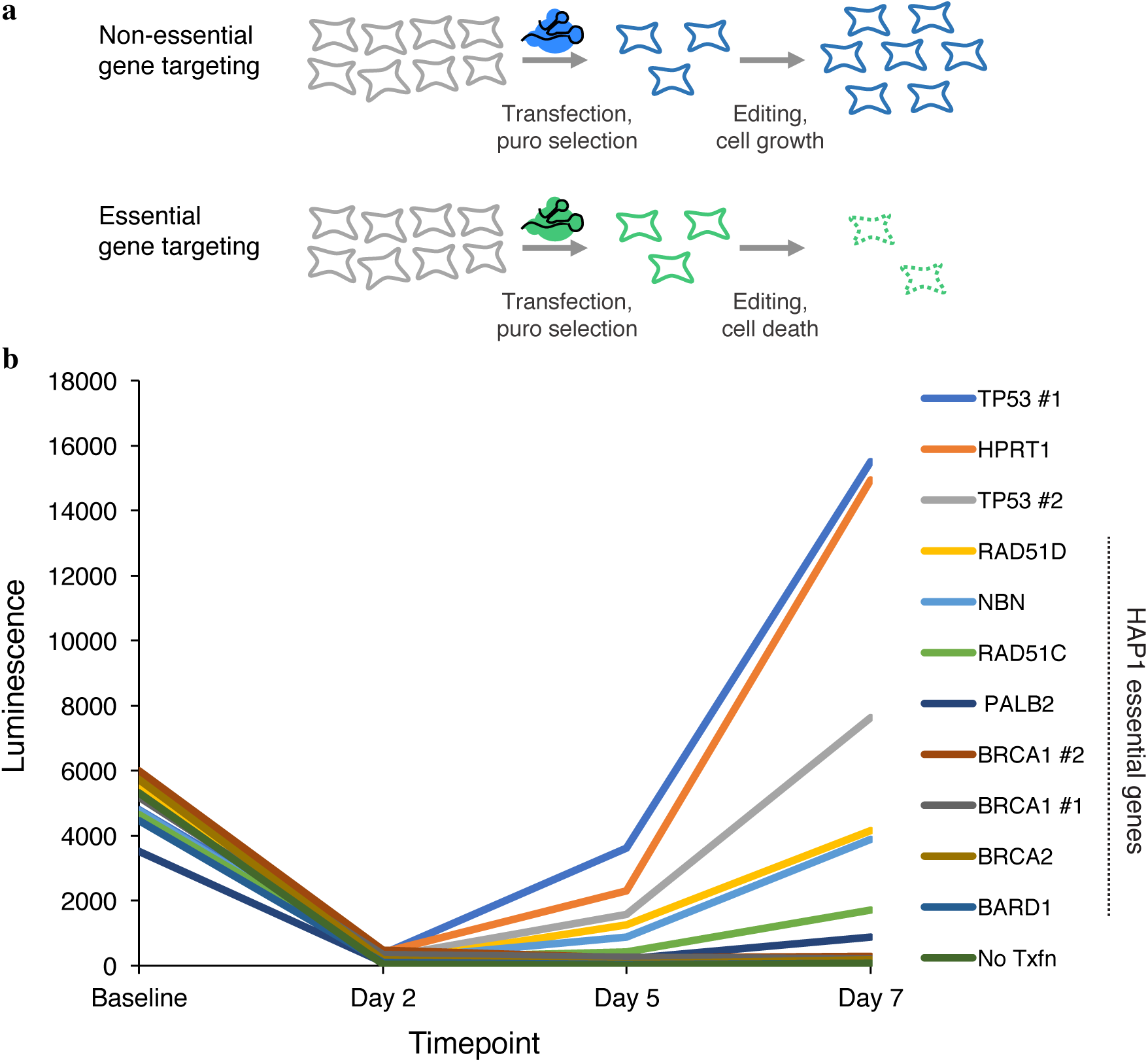
CRISPR targeting of HDR pathway genes to confirm essentiality in HAP1 cells. **a**, Schematic; HAP1 cells are transfected with a plasmid expressing a gRNA and a Cas9-2A-puromycin cassette^29^. Due to low transfection rates for HAP1 cells, puromycin selection reduces viable cells in all transfections. Over time, however, CRISPR targeting of non-essential genes leads to increased cell growth compared to CRISPR targeting of essential genes. **b**, Cell viability of HAP1 cells transfected with Cas9/gRNA constructs targeting different HDR genes and controls (*HPRT1*, *TP53*) was measured using the CellTiterGlow assay. Luminescence is proportional to the number of living cells in each well when the assay is performed. Triplicate wells for each gRNA at each time point were processed, quantified on a plate reader and averaged. gRNA sequences are included in Supplementary Table 2.

**Extended Data Figure 2.**
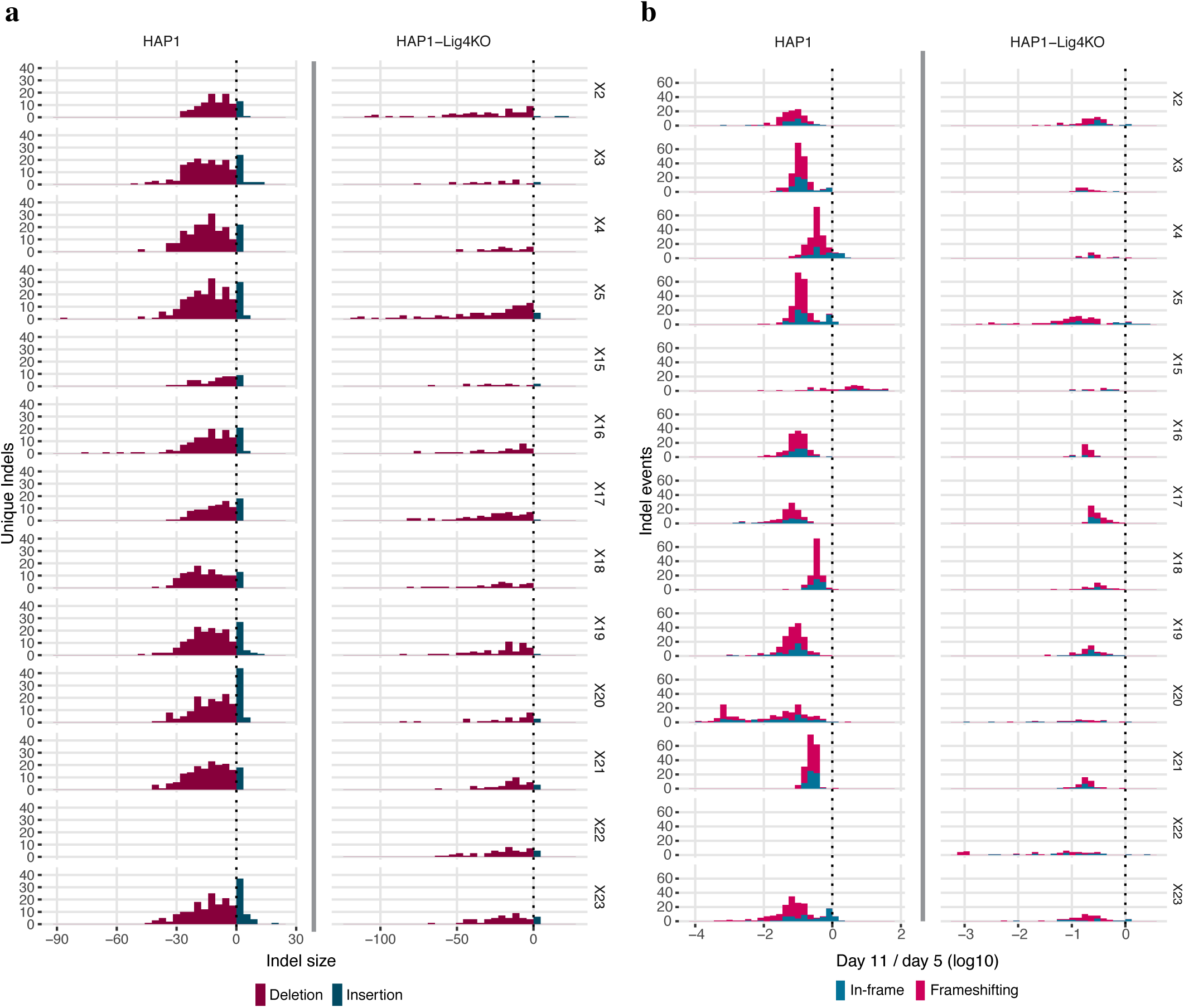
Analysis of Cas9-induced indels observed in *BRCA1* SGE experiments. Variants observed in gDNA sequencing were included in this analysis if i) they aligned to the reference with either a single insertion or deletion within 15 bp of the predicted Cas9 cleavage site and ii) were observed at a frequency greater than 1 in 10,000 reads in both replicates. **a**, Histograms show the number of unique indels observed of each size, with negative sizes corresponding to deletions. More unique indels were observed in WT HAP1 cells compared to HAP1-Lig4KO cells for exons compared (WT data for exon 22 was excluded). **b**, Day 11 over day 5 indel frequencies were normalized to the median synonymous SNV in each replicate and then averaged across replicates to measure selection on each indel. The distribution of selective effects is shown for each experiment as a histogram, in which indels are colored by whether their size was divisible by 3 (*i.e.* ‘in-frame’ vs. ‘frameshifting’). Whereas frameshifting variants were consistently depleted, some exons were tolerant to in-frame indels.

**Extended Data Figure 3.**
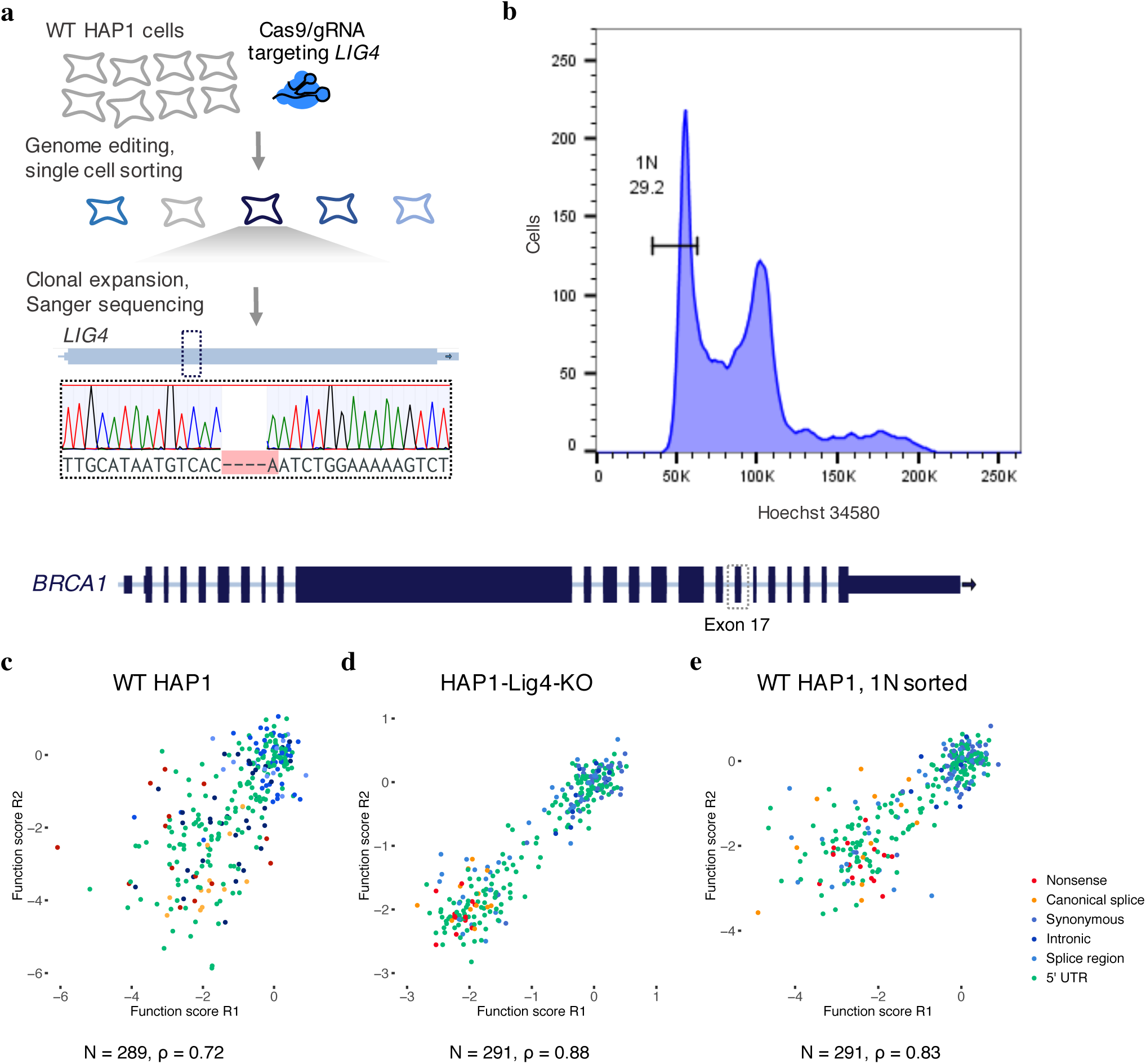
HAP1 cell line optimizations for saturation genome editing to assay essential genes. **a**, A gRNA targeting Cas9 to the coding sequence of *LIG4*, a gene integral to the non-homologous end-joining pathway, was cloned into a vector co-expressing Cas9-2A-GFP^29^. WT HAP1 cells were transfected, and single GFP-expressing cells were sorted into wells of a 96-well plate. Eight monoclonal lines were grown out over a period of three weeks and screened using Sanger sequencing for frameshifting indels in *LIG4*. The Sanger trace shows the frameshifting deletion present in the clonal line chosen for subsequent experiments, referred to as ‘HAP1-Lig4KO’. **b**, To purify HAP1 cells for haploid cells, live cells were stained for DNA content with Hoechst 34580 and sorted using a gate to select cells with the lowest DNA content, corresponding to 1N cells in G1. **c-e**, Plots comparing SNV function scores across replicate experiments for exon 17 saturation genome editing experiments performed in unsorted WT HAP1 cells (**c**), HAP1-Lig4KO cells (**d**), and WT HAP1 cells sorted on 1N ploidy (**e**). Both *LIG4* knockout and 1N-sorting improved replicate correlations.

**Extended Data Figure 4.**
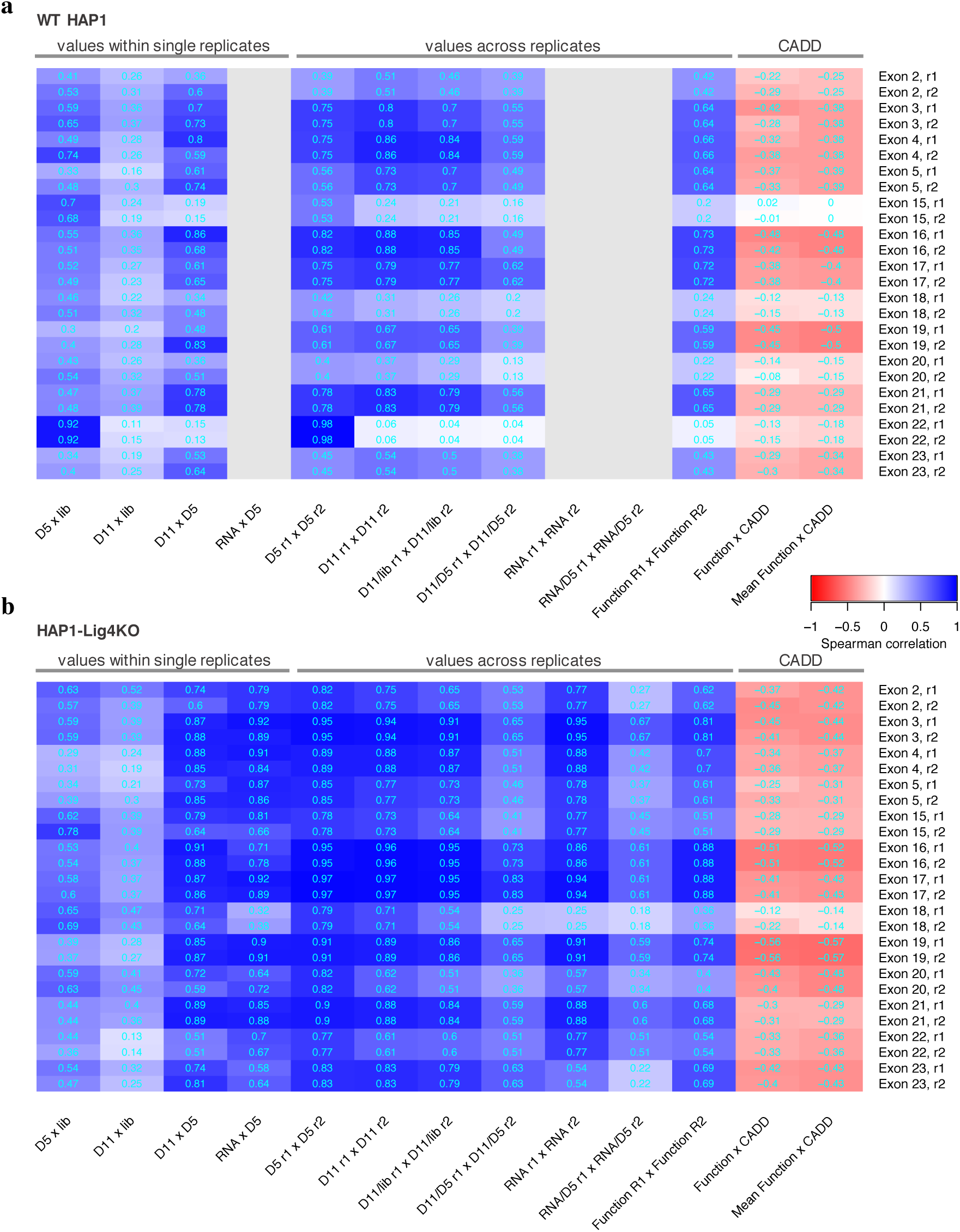
Correlations for SNV measurements within single experiments, across transfection replicates, and to CADD scores for all SGE experiments. Heatmaps indicate Spearman correlation coefficients for SNV measurements from experiments in WT HAP1 cells (**a**) and in HAP1-Lig4KO cells (**b**). Gray boxes indicate absent RNA data from WT HAP1 cells. The four leftmost columns show how SNV frequencies correlate between samples from within a single replicate experiment. The unusually high correlations between exon 22 SNV frequencies in the plasmid library and in day 5 gDNA samples from WT HAP1 cells suggests plasmid contamination in gDNA. Indeed, primer homology to a repetitive element in the exon 22 library was identified. Consequently, the WT HAP1 exon 22 data was removed from analysis and a different primer specific to gDNA was used to prepare exon 22 sequencing amplicons from HAP1-Lig4KO cells. The low HAP1-Lig4KO correlations between exon 18 SNV frequencies in day 5 gDNA and RNA and between RNA replicates suggests RNA sample bottlenecking consequential to low RNA yields. Therefore, exon 18 RNA was also excluded from analysis. Consistent with the higher rates of HDR-mediated genome editing (**Fig. 2a**), replicate correlations (middle columns) were generally higher in HAP1-Lig4KO cells than WT HAP1 cells. CADD scores predict the deleteriousness of each SNV, and are therefore negatively correlated with function scores (rightmost columns).

**Extended Data Figure 5.**
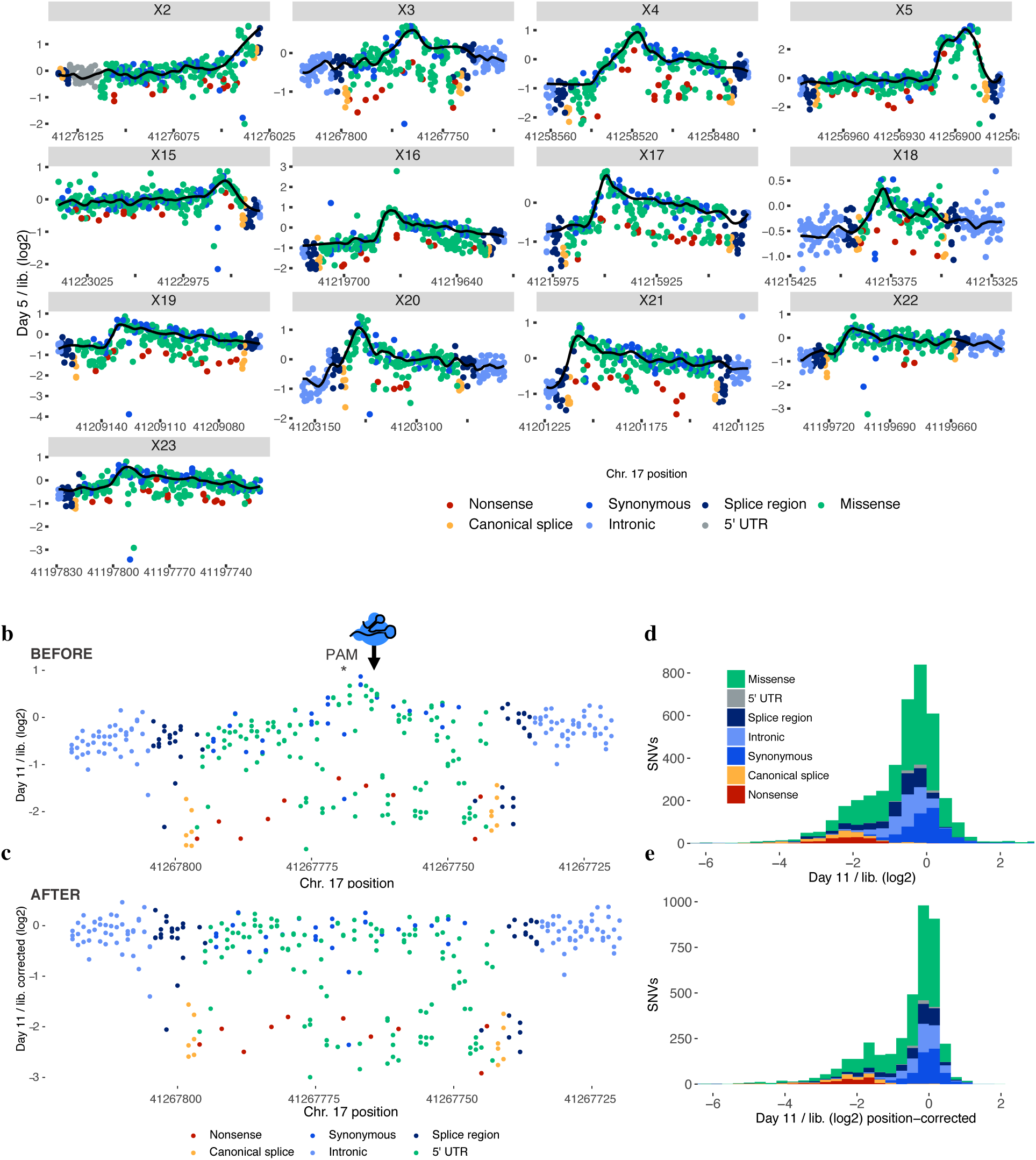
Models of SNV editing rates across *BRCA1* exons account for positional biases. **a**, Gene conversion tracts arising during HDR in human cells are short such that library SNVs are introduced to the genome more frequently near the CRISPR target site. We modelled this positional effect in our data using a LOESS regression fit on day 5 over library SNV ratios. Plots shown here are of the average of two replicate experiments per exon, with the black line indicating the LOESS regression. By day 5 sampling, selective effects on gene function are evidenced by nonsense SNVs (red) appearing at lower frequencies compared to neighbouring SNVs. Therefore, to best approximate the SNV editing rate as a function of position alone (*i.e.* the ‘baseline’), the regression excluded SNVs that were selected against between day 11 and day 5 (see Methods). **b**,**c**, Day 11 over library SNV ratios were adjusted by the positional fit for each experiment in calculating function scores. This adjustment is illustrated here for an exon 3 replicate by plotting the ratio as a function of position before (**b**) and after (**c**) adjustment. The elevated day 11 over library ratios for SNVs near the CRISPR target site are corrected to achieve a more uniform baseline across the mutagenized region. **d**,**e**, The distributions of SNV day 11 over library ratios before and after accounting for positional effects are shown, colored by mutational consequence (pre-filtering, N = 4,002).

**Extended Data Figure 6.**
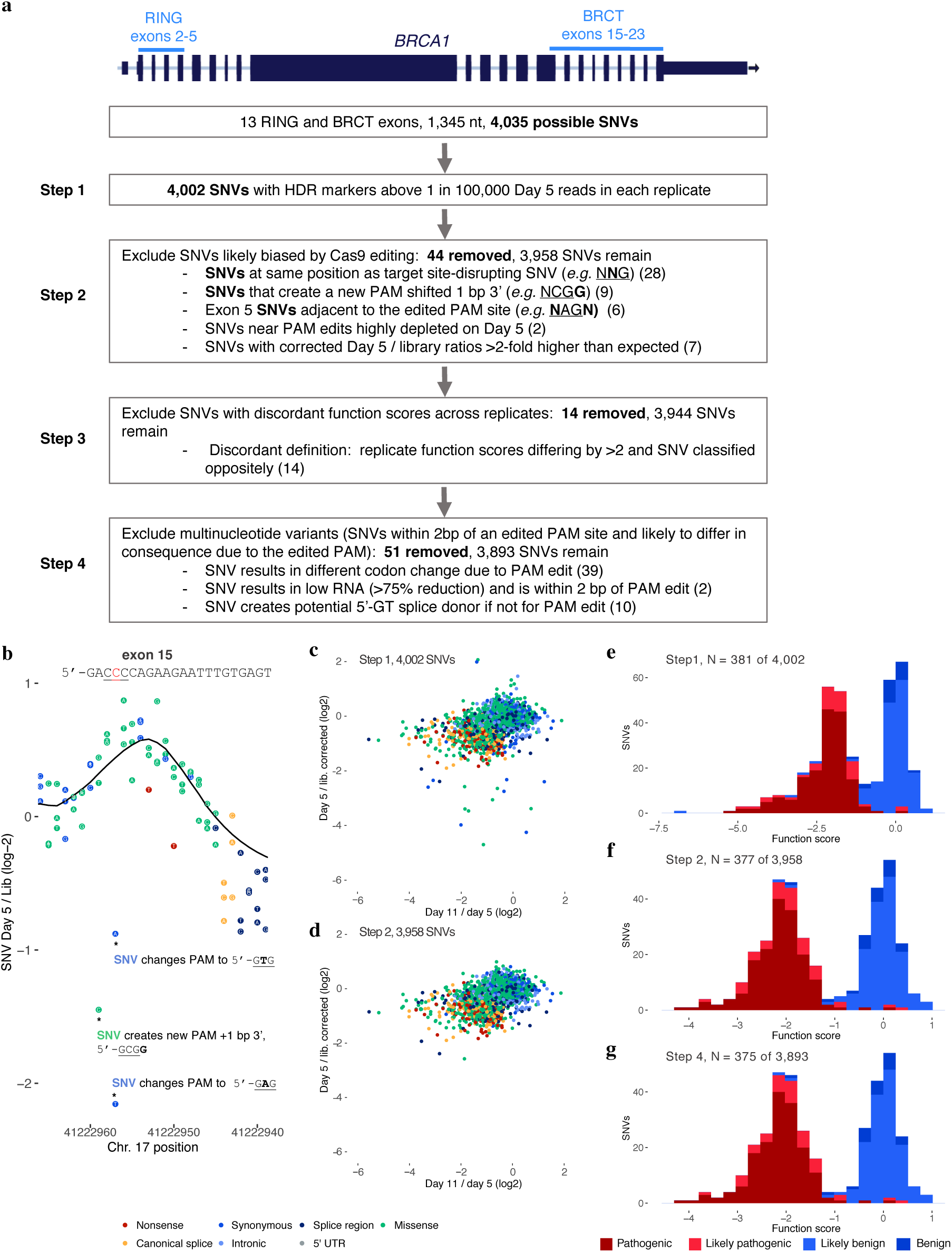
SNV filtering to prevent erroneous functional classification. **a**, The flow chart describes filters used to produce the final SNV data set and shows how many SNVs were removed at each step. **b**, Raw day 5 over library SNV ratios are shown for a portion of exon 15 to illustrate how re-editing biases necessitate filtering. The three depleted SNVs marked with asterisks create alternative PAM sequences that likely allow the Cas9:gRNA complex to re-cut the locus and cause their removal. For other SNVs, the fixed PAM edit (a <u>GGG</u> to <u>GCG</u> synonymous change) minimalizes re-editing. The location of the target PAM is underlined and each indicated SNV is bolded in the annotations. The LOESS regression curve in shown in black. **c**,**d**, Plots show the relationship between day 5 over library and day 11 over day 5 ratios before (**c**) and after (**d**) filtering steps 1 and 2. Filtering removes outliers because editing biases primarily affect the day 5 over library ratio. **e-g**, Histograms show the distributions of function scores for SNVs deemed ‘pathogenic’ or ‘benign’ in ClinVar at different stages of filtering. Scores in **e** are derived prior to normalization across exons.

**Extended Data Figure 7.**
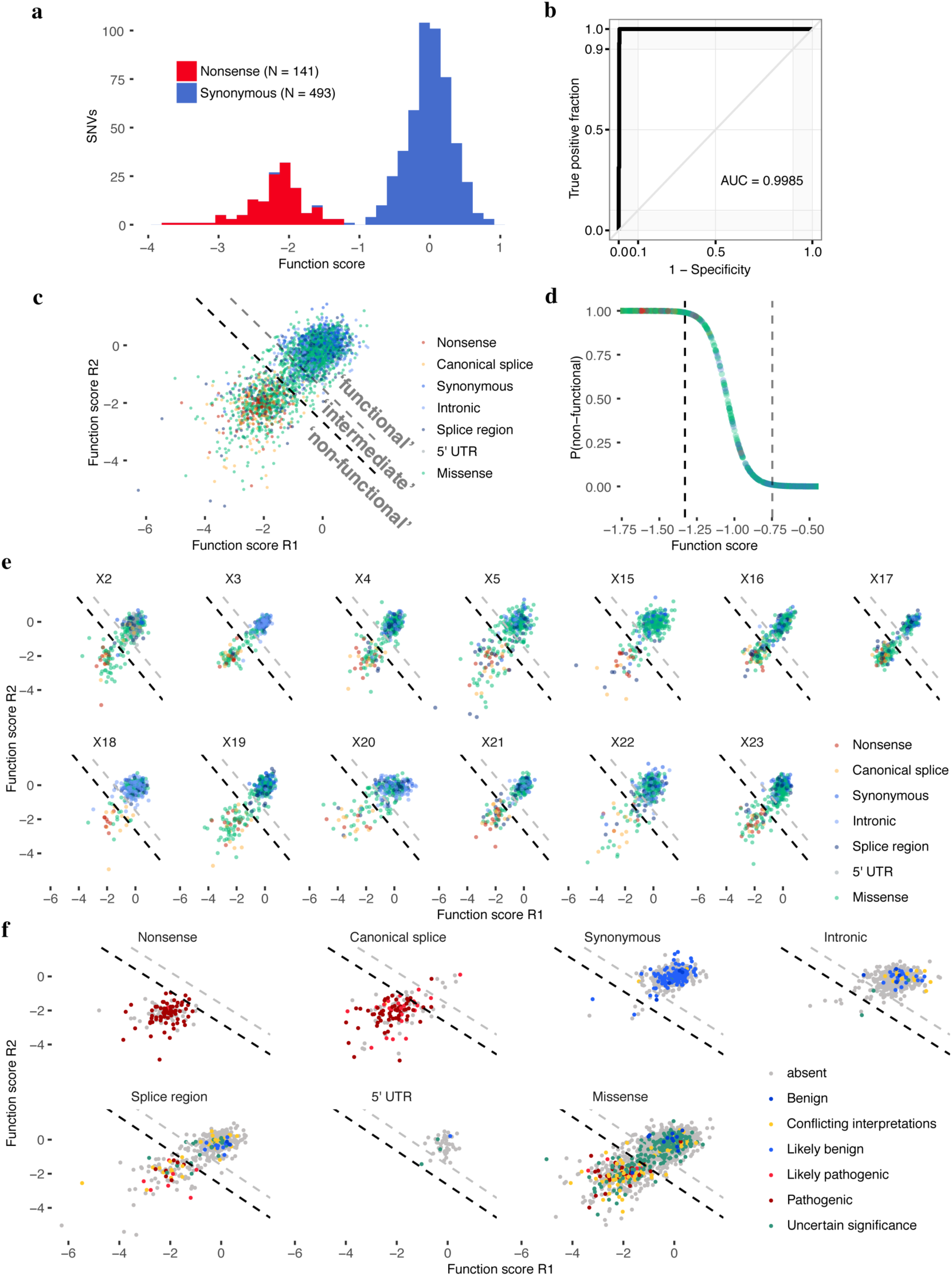
Mixture modeling of scores to classify SNVs by functional effect. **a**, Distributions of ‘non-functional’ and ‘functional’ SNVs plotted here were defined respectively as all nonsense SNVs and all synonymous SNVs with RNA scores within 1 SD of the median synonymous SNV. **b**, An ROC curve was generated using SGE function scores to distinguish the 634 ‘functional’ and ‘non-functional’ SNVs defined in **a**. **c**, A two-component Gaussian mixture model was used to produce point estimates of the probability that each SNV was ‘non-functional’, *P*(nf), given its average function score across replicates. These P-values are plotted in **d** against function scores for a subset of the data. Thresholds were set such that *P*(nf) < 0.01 corresponds to ‘functional’, and *P*(nf) > 0.99 corresponds to ‘non-functional’, and 0.01 < *P*(nf) < 0.99 corresponds ‘intermediate’ classification. Functional classification thresholds are drawn as dashed lines; black denotes the non-functional threshold and gray the intermediate threshold. **e**,**f**, SNV function scores across replicates are plotted for each exon with SNVs colored by mutational consequence (**e**), and for each type of mutational consequence with SNVs colored by ClinVar status (**f**). Using the optimal function score cutoff for all SNVs tested (Fig. 3b), sensitivities and specificities for distinguishing ‘Pathogenic’/’Likely pathogenic’ from ‘Benign’/’Likely benign’ ClinVar annotations for each type of mutation are as follows: 92.7% and 92.9% for missense SNVs (N = 55), 100% and 100% for splice region SNVs (N = 23), and 95.2% sensitivity for canonical splice site SNVs (N = 83; specificity not calculable).

**Extended Data Figure 8.**
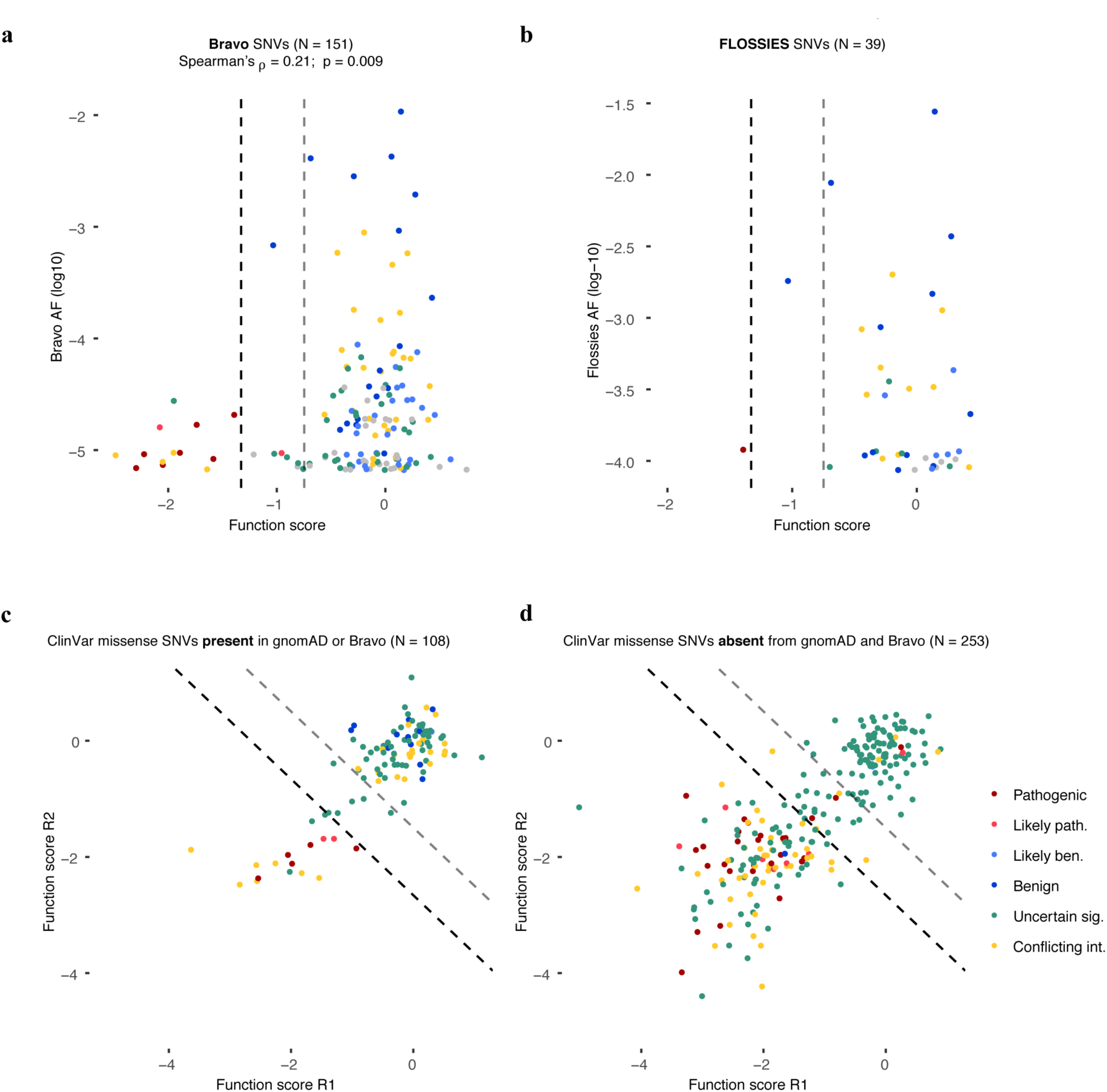
*BRCA1* SNVs observed more frequently in large-scale population sequencing are more likely to score as functional. SNV function scores are plotted against Bravo allele frequencies (**a**) and FLOSSIES allele frequencies (**b**). **a**, Bravo is a collection of whole genome sequences ascertained from 62,784 individuals through the NHLBI TOPMed program. Similarly to SNVs present in gnomAD (Fig. 3d), higher allele frequencies of SNVs in Bravo correlate with higher function scores. **b**, FLOSSIES is a database of variants seen in targeted sequencing of breast cancer genes sampled from approximately 10,000 cancer-free women at least 70 years old. Only 1 of 39 SNVs observed in FLOSSIES scored as non-functional. **c**,**d**, Missense SNVs in ClinVar are separated by whether they have (**c**) or have not (**d**) been seen in either gnomAD or Bravo and function scores across replicates are plotted, with dashed lines demarcating functional classes. A higher proportion of ClinVar missense SNVs absent from gnomAD and Bravo score as non-functional (50.6% vs. 15.7%, Fisher’s exact *P* = 1.80 × 10^−17^).

**Extended Data Figure 9.**
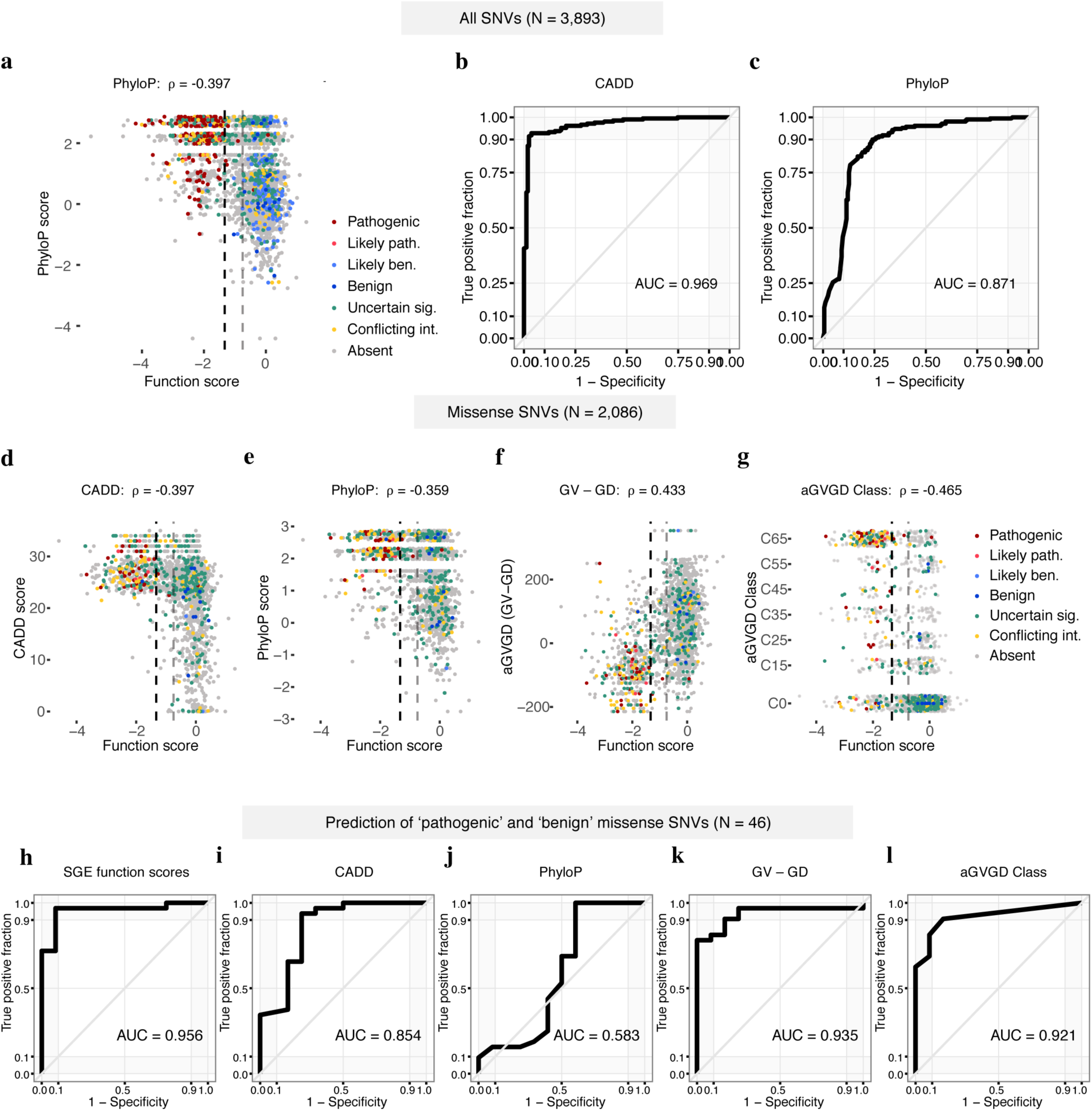
SGE function scores correlate with computational metrics and perform favorably at predicting ClinVar annotations. **a**, SNV function scores are plotted against mammalian phyloP scores, with colors indicative of ClinVar status. **b**,**c**, ROC curves show the performance of CADD scores and phyloP scores for discriminating ClinVar ‘pathogenic’ and ‘benign’ SNVs (including ‘likely’), as described in Fig. 3b for SGE data. **d-g** Plots as in **a**, but for missense SNVs only, showing correlations between SGE function scores and CADD^39^ scores, phyloP scores^40^, Grantham differences (Grantham amino acid variation minus Grantham amino acid deviation; GV - GD), and align-GVGD classifications^53^. Missense SNV function scores also correlate with SIFT scores^54^ (ρ = 0.363) and PolyPhen-2 scores^55^ (ρ = −0.277). (*P* < 1 × 10^−37^ for all correlations.) **h-l**, ROC curves assess the performance of SGE function scores and each indicated metric at distinguishing firmly ‘pathogenic’ and ‘benign’ missense SNVs. (*i.e.* not including ‘likely’).

**Extended Data Figure 10.**
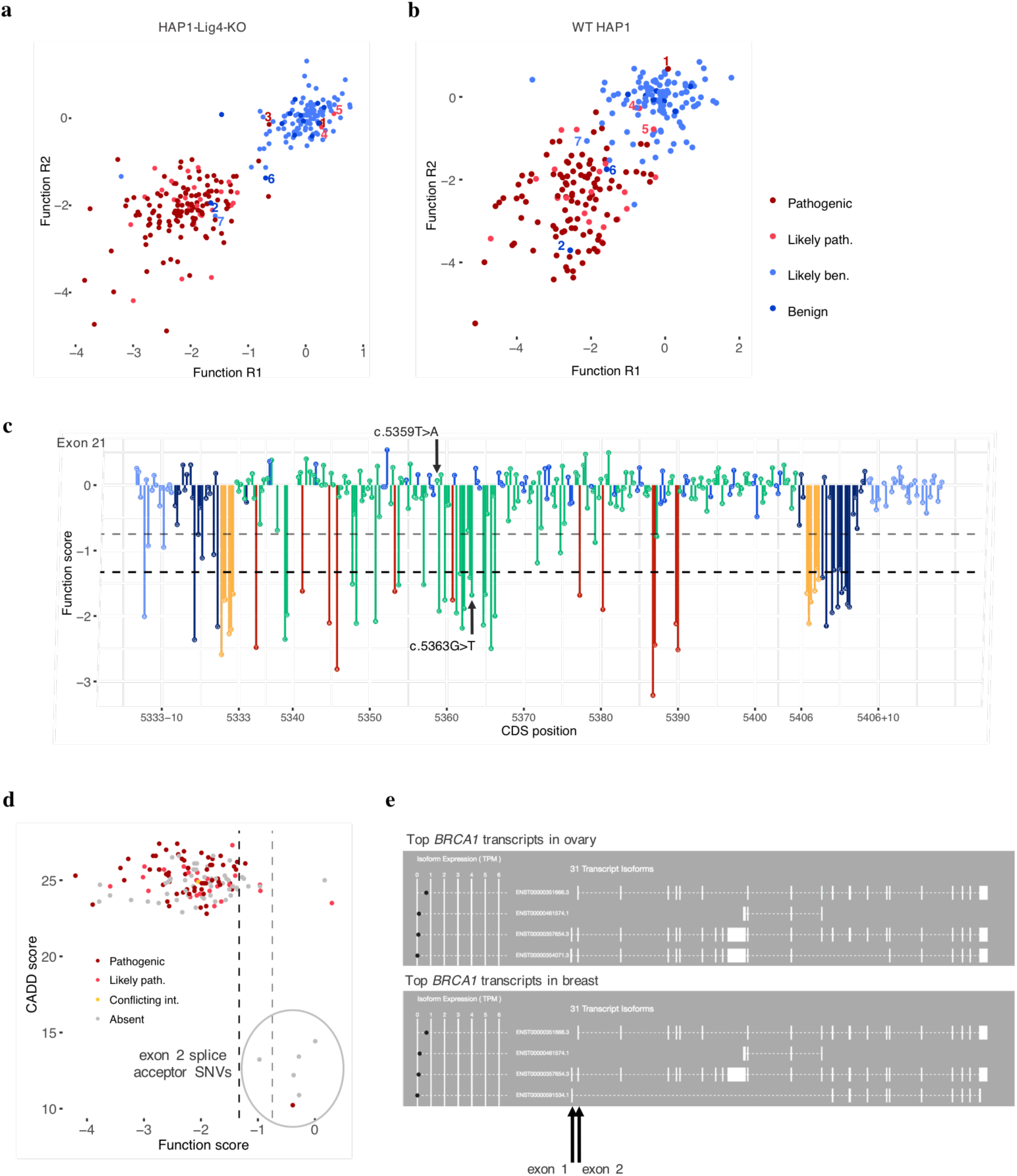
Evidence supporting SNV scores in discordance with ClinVar classifications. Function scores of SNVs classified as ‘benign’ or ‘pathogenic’ (including likely’s) are shown across replicates for experiments using HAP1-Lig4KO cells (**a**) and for preliminary experiments using WT HAP1 cells (**b**). Plots exclude exons with low overall reproducibility in WT HAP1 cells (replicate correlations < 0.4: exons 15, 18, 20 and 22). The three SNVs firmly discordant with ClinVar are labelled 1-3 in **a**, corresponding to c.5359T>A (dark red 1), c.5044G>A (dark blue 2), and c.-19-2A>G (dark red 3), respectively. The same filtering criteria were applied to both sets of experiments, which led to the removal of SNV 3 from the WT HAP1 data due to disagreement of scores between replicates. Discordant ‘likely pathogenic’ SNVs (4,5), an intermediate scoring ‘benign’ SNV (6) and a discordant ‘likely benign’ SNV (7) are also labelled for comparison. **c**, The sequence-function map of exon 21 is shown with the function scores for the two ‘pathogenic’ SNVs observed in linkage indicated. Dashed lines demarcate functional classifications. **d**, Function scores are plotted against CADD scores for all canonical splice SNVs assayed, colored by ClinVar status. The six possible exon 2 splice acceptor SNVs (circled) have the lowest CADD scores among all canonical splice SNVs assayed, and none score as ‘non-functional’. **e**, GTEx browser shots show that many of the most common *BRCA1* transcripts mapped from ovarian and breast tissues lack the exon 1 / exon 2 junction.

